# VR-based real-time imaging reveals abnormal cortical dynamics during behavioral transitions in a mouse model of autism

**DOI:** 10.1101/2022.11.14.516121

**Authors:** Nobuhiro Nakai, Masaaki Sato, Okito Yamashita, Yukiko Sekine, Xiaochen Fu, Junichi Nakai, Andrew Zalesky, Toru Takumi

## Abstract

Functional connectivity (FC) can provide insight into cortical circuit dysfunction in neuropsychiatric disorders. However, dynamic changes in FC related to locomotion with sensory feedback remain unexplored. To investigate FC dynamics in locomoting mice, we developed mesoscopic Ca^2+^ imaging with a virtual reality (VR) environment. We find rapid reorganization of cortical FC in response to changing behavioral states. Using machine learning classification, behavioral states are accurately decoded. We then use our VR-based imaging system to study cortical FC in a mouse model of autism and find that locomotion states are associated with altered FC dynamics. Furthermore, we identify FC patterns involving the motor area as the most distinguishing features of the autism mice from wild-type mice during behavioral transitions, which might correlate with motor clumsiness in patients with autism. Our VR-based real-time imaging system provides invaluable information to understand FC dynamics linked to a behavioral abnormality of neuropsychiatric disorders.

## Introduction

Neocortical activity displays dynamic changes across multiple cortical areas to facilitate processing of sensory information and generate action outputs ^1^. Such large-scale network dynamics can be investigated using functional connectivity (FC), defined as temporal dependence of neuronal activity between anatomically separated brain regions ^2^. With functional magnetic resonance imaging (fMRI), FC is quantified as the extent of coactivation between spontaneous blood-oxygen-level-dependent (BOLD) signals during rest ^2,3^ or during task conditions necessitating minimal movement. FC can also be measured in rodents ^4^. However, immobilization of a subject within an MRI scanner and the slow nature of BOLD signals in resting-state fMRI has limited the study of cortical activity during complex behaviors involving whole-body movement and locomotion. Although the ability to record sensory-evoked BOLD signals in awake head-restrained mice was recently developed ^5^, techniques to measure FC during natural and voluntary movement in an interactive environment remain to be established.

FC provides a valuable tool for investigating functional brain network organization in autism spectrum disorder (ASD) ^6^. A large body of resting-state fMRI studies reports functional under-connectivity (hypo-connectivity), over-connectivity (hyper-connectivity), and a combination of both global and local alterations in the ASD brain ^7^. In addition, machine learning models can be trained to predict an individual’s diagnostic status using their FC, although clinical heterogeneity is a significant challenge ^8,9^. In contrast to resting-state and task conditions, cortical dynamics during voluntary behaviors such as locomotion remain to be understood, particularly in neuropsychiatric disorders. Individuals with ASD exhibit motor coordination deficits ^10^ and impairment of movement planning in goal-directed locomotion ^11,12^. Furthermore, accumulating evidence suggests that sensorimotor difficulties seen in ASD are strongly associated with the development and maintenance of social and non-social core symptoms ^13^.

In this study, we sought to elucidate the rapid reorganization of functional cortical networks during locomotion, focusing on periods transitioning between locomotion (i.e., running) and rest conditions, in normal and ASD model mice. To this end, we developed an integrated Ca^2+^ imaging and virtual reality (VR) platform to study neural activity in mice during VR locomotion, including statistical analysis of second-by-second FC dynamics, graph theoretical analysis of network structures, and machine learning classification of FC patterns using support vector machine (SVM). Cortex-wide mesoscopic Ca^2+^ imaging enabled the measurement of neural activity with high spatiotemporal resolution ^14,15^. VR created an environment that simulated real-world situations for head-fixed mice and allowed us to manipulate sensory information ^16,17^. Using this experimental and analytical framework, we assessed cortical FC of a copy number variation mouse model for human 15q11-13 duplication (*15 dup*) in different behavioral states. We previously reported that *15q dup* mice display ASD-like social communication deficits ^18,19^ and exhibit abnormal somatosensory tuning under anesthesia and whole-brain functional hypoconnectivity in awake resting-state fMRI ^5,20^. However, cortical FC alterations during behavior remain unknown. Here, we found that these mice exhibited impaired locomotion-dependent FC dynamics and aberrant FC patterns involving hyperconnectivity of the motor areas, highlighting the importance of motor areas in cortical FC dysfunction during spontaneous behavioral switching in ASD.

## Results

To measure cortical FC in mice engaged in voluntary movement, we used transcranial Ca^2+^ imaging combined with a head-fixed VR system (**Figures 1A–1C; STAR Methods**). The virtual environment mimicked a realistic open-field enclosure and consisted of a two-dimensional square arena with differently colored walls (**Figure 1B**). We crossed Emx1-Cre driver mice, which allows extensive Cre-mediated recombination in the forebrain, with Ai95D mice to express the genetically-encoded calcium indicator GCaMP6f in cortical excitatory neurons of the offspring Emx1G6 mice (**Figure 1D; STAR Methods**). During a 10-min VR session, mice exhibited voluntary locomotion (speed > 0.5 cm/s) in this virtual arena; they spent 58.7 ± 19.7 % of the total duration in a state of locomotion (mean ± SD, *n* = 89 sessions from 7 mice). For further analysis, we excluded periods of frequent alterations between locomotion and rest and focused only on long episodes (continuously ≥ 3 s) of locomotion and rest (**Figure 1E**). The percentage of time spent in long locomotion and rest was 50.3 ± 22.1 % and 32.5 ± 20.2 %, respectively (mean ± SD, *n* = 89 sessions from 7 mice). Average lengths of long locomotion and rest episodes were 10.0 ± 5.4 s (mean ± SD, *n* = 2,836 episodes from 89 sessions) and 9.4 ± 6.6 s, (mean ± SD, *n* = 1,903 episodes from 89 sessions), respectively. There were no significant changes in these behavioral parameters across sessions (**Figure 1F**). We imaged cortical fluorescence changes at a frame rate of 30 frames per second in 50 ROIs (regions of interest) that covered most of the dorsal cortical subregions (**Figures 1G and 1H; Figures S1 and S2; STAR Methods**). Pair-wise correlation coefficients were computed between cortical ROIs at a temporal scale of a second, using a one-frame sliding window. We then applied graph-theoretic analyses to characterize the resulting network dynamics and visualized highly correlated ROI pairs (*r* > 0.8) using an FC map (**Figure 1H; STAR Methods**).

**Figure 1.**
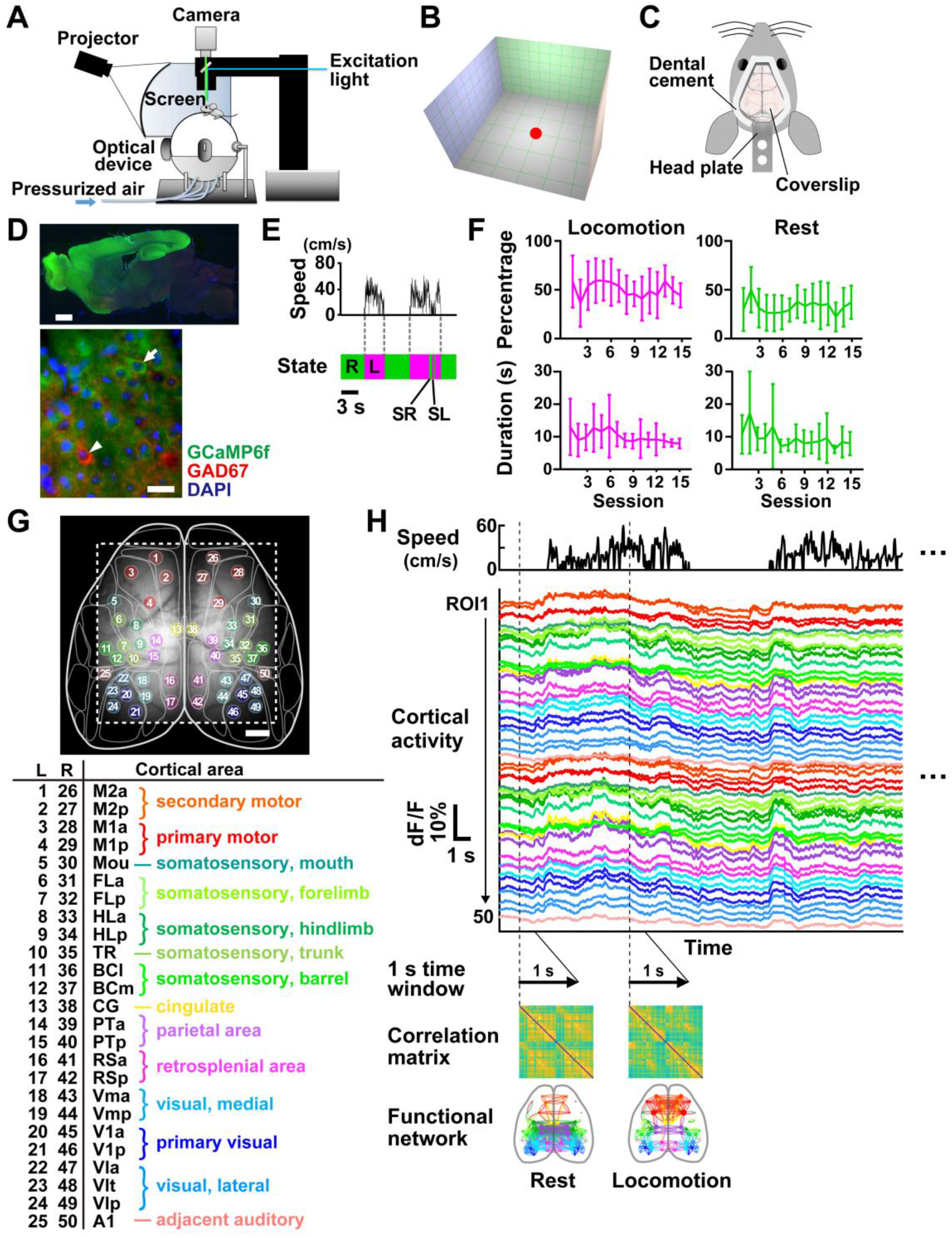
Analysis of cortical functional connectivity with mesoscopic Ca^2+^ imaging. **(A)** The imaging and virtual reality (VR) system. **(B)** The virtual arena. The floor and walls have green gridlines to enhance the sense of visual flow. Each wall is painted in a different color. The mouse starts to move from the location indicated by the red dot. **(C)** A schematic of transcranial imaging window affixed to the mouse skull. **(D)** Expression of GCaMP6f in a parasagittal section of an adult Emx1G6 mouse (top, scale bar = 1 mm). Immunofluorescence detection of GCaMP6f (green) and GAD67 (red) in layer 2/3 of the primary motor cortex (bottom, scale bar = 20 μm). Cell nuclei were stained with DAPI (blue). The arrow and arrowhead indicate an example of GCaMP6f-positive and GAD67-positive cells, respectively. **(E)** Two behavioral states, long locomotion (L) and long rest (R), were defined by spontaneous locomotion and resting states (duration: ≥3 s) of head-fixed mice. Locomotion and rest episodes shorter than 3 s, short locomotion (SL) and short rest (SR) were excluded from functional connectivity analysis during behavioral transitions. **(F)** Percentages of time spent in long locomotion and long rest (top) and average lengths of long locomotion and long rest episodes (bottom) across sessions. Data represent mean ± SD. (Percentage) locomotion: *F*_(14, 74)_ = 0.61, *P* = 0.84; rest: *F*_(14, 74)_ = 0.52, *P* = 0.91; (Duration) locomotion: *F*_(14, 74)_ = 0.63, *P* = 0.83; rest: *F*_(14, 74)_ = 1.83, *P* = 0.58, *n* = 7 mice, one-way ANOVA. **(G)** Fifty cortical ROIs are overlaid onto a grayscale image of the dorsal cortex with a cortical parcellation map (top, dashed lines indicate the field of view, scale bar = 1 mm). ROIs 1–25 and 26–50 were defined in the left (L) and right (R) hemispheres, respectively, and ROIs for each hemisphere were numbered along the anterior-posterior axis (bottom). The lower case letters following cortical areas indicate anterior (e.g., M2a) and posterior (e.g., M2p), or lateral (e.g., BCl) and medial (e.g., BCm) positions. **(H)** Analysis of cortical functional connectivity. After calculating normalized fluorescence changes (dF/F) for each ROI, pair-wise Pearson’s correction coefficients of cortical activity in a one-second time window were calculated for all ROI pairs and then visualized as matrices. Each matrix was labeled with a corresponding behavior state at the first frame of the time window. In graph visualization of functional networks, connectivity with a correlation coefficient above a threshold (*r* > 0.8) was denoted as a line (edge) that connected the corresponding ROIs (nodes).

### Graph analysis of cortical network dynamics during behavioral transitions

First, we examined the activity of different cortical areas during behavioral transitions from long rest to long locomotion (locomotion onset, *n* = 566 events from 89 sessions) and from long locomotion to long rest (locomotion cessation, *n* = 643 events from 89 sessions; **Figure 2A**). During a period that spanned 3 s before and after the onset of locomotion, many cortical areas displayed marked transient increases in fluorescence intensity that began slightly prior to the onset (dF/F; 0.05 ± 0.27 % at −0.5 s; 0.62 ± 0.45 % at 0 s; *n* = all 50 ROIs, mean ± SD; **Figure 2A**). In contrast, the fluorescence intensity of all areas rapidly and substantially decreased immediately after the cessation of locomotion (−0.06 ± 0.12 % at 0 s; −0.54 ± 0.34 % at 0.5 s; **Figure 2A**). Such large signal changes were not observed during control periods that were randomly selected independent of the locomotion state (−0.01 ± 0.03 % at −0.5 s; −0.03 ± 0.04 % at 0 s; 0.01 ± 0.03 % at 0.5 s; **Figure 2A**). Hierarchical clustering of regional fluorescence signals revealed that response profiles of all ROIs during the onset periods could be divided into two major clusters; one represented the considerable transient activity of sensory (V1, HL, FL, etc.) and association (PT, RS, etc.) areas and the other represented sustained activity of the motor-related regions (M1, M2, etc.; **Figure S3A**). On the other hand, clustering of activity around the locomotion cessation differentiated M2 and CG from the major clusters (**Figure S3B**).

**Figure 2.**
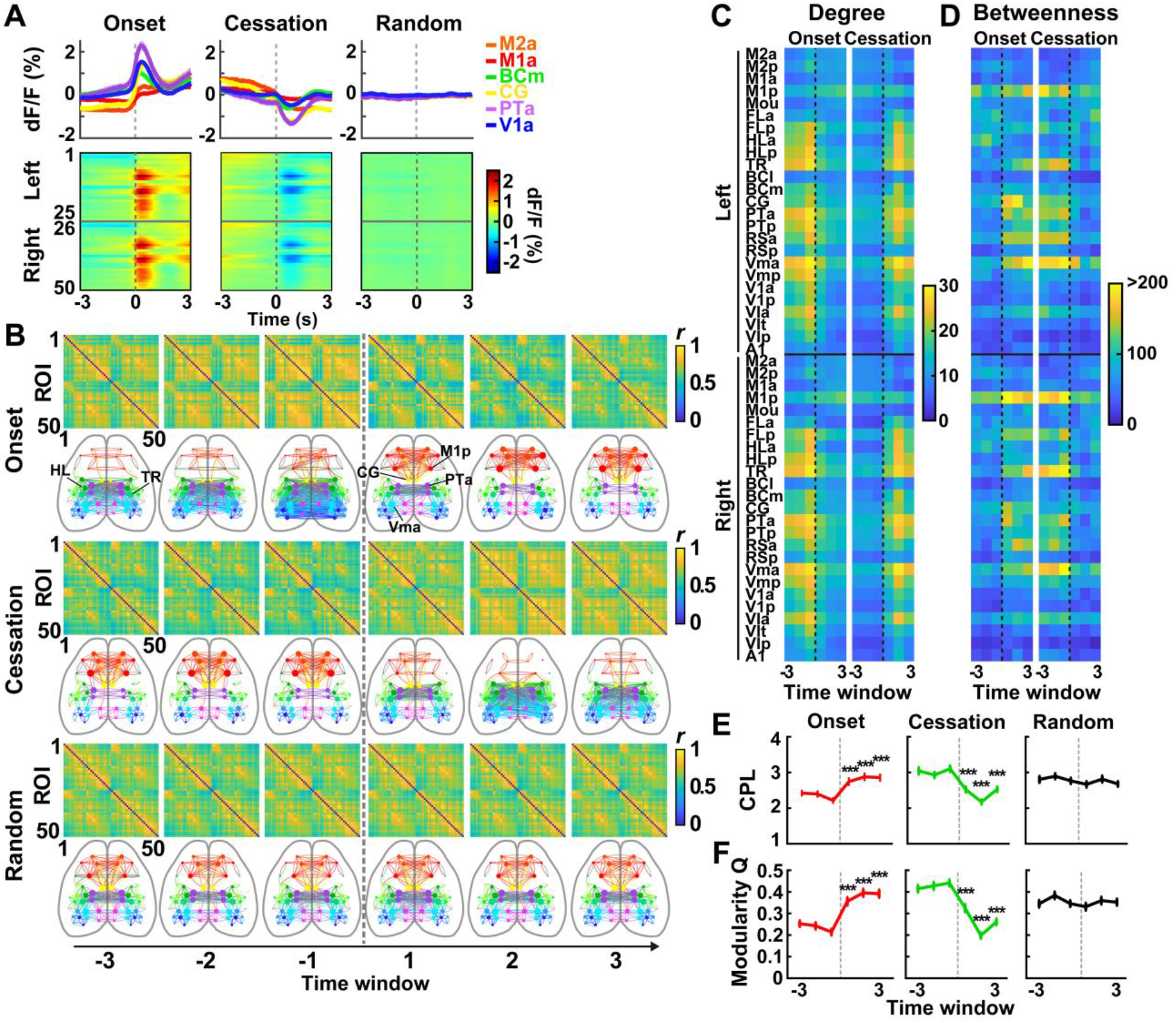
Dynamic reconfiguration of the functional cortical network during behavioral transitions. **(A)** Cortical activity during behavioral transitions in Emx1G6 mice. The top plots present average relative changes in fluorescence signals in representative cortical areas (*n* = 89 sessions from 7 mice). The vertical dashed lines indicate the occurrence of transition. The colormaps at the bottom show changes in fluorescence signals in all ROIs. ROI 1–25 and 26–50 were defined in the left and right hemispheres, respectively (see Figure 1G for details). **(B)** Dynamics of the functional cortical network during the behavioral transitions. The data for locomotion onset, cessation, and random control are shown from top to bottom. Correlation matrices and functional connectivity graphs (FC, *r* > 0.8) were determined for each second of the time window encompassing the relevant behavioral transition that occurred at time zero (vertical dashed line). Time windows -3, -2, -1, 1, 2, 3, correspond to windows that cover -3 s to -2 s, -2 s to -1 s, -1 s to 0 s, 0 s to 1 s, 1 s to 2 s and 2 s to 3s, respectively. (**C, D**) Changes in node degree (C) and betweenness centrality (D) during the transitions. **(E)** Change in characteristic path length (CPL) during the transitions. Data represent mean ± SEM. Onset: *F*_(5, 612)_ = 11.35, *P* = 1.7×10^−10^, Cessation: *F*_(5, 600)_ = 19.15, *P* = 1.1×10^−17^, Random: *F*_(5, 612)_ = 0.22, *P* = 0.95, one-way ANOVA. \*\*\**P* < 0.001, vs. time window -1, Tukey Kramer test, *n* = 89 sessions from 7 mice. **(F)** Change in modularity Q during the transitions. Data represent mean ± SEM., Onset: *F*_(5, 612)_ = 25.46, *P* = 2.3×10^−23^, Cessation: *F*_(5, 600)_ = 37.20, *P* = 3.0×10^−23^, Random: *F*_(5, 612)_ = 0.51, *P* = 0.77, one-way ANOVA. \*\*\**P* < 0.001, vs. time window -1, Tukey Kramer test, *n* = 89 sessions from 7 mice.

Next, we investigated cortical FC dynamics during transition periods by visualizing indices that represent network centrality of each cortical area. Node degree captures the extent to which a region connects with other regions. Betweenness centrality measures how much a region is in-between other regions ^21^. Before locomotion onset, FC among posterior areas (blue and green edges), most notably bilateral HL, TR, PT, and Vm, gradually increased (time window from −3 to −1, **Figure 2B top**) and node degree also increased in many areas (15.8 ± 5.9 at −3 s; 20.9 ± 6.3 at −1 s; *n* = all 50 ROIs, mean ± SD; **Figure 2C**). At locomotion onset, FC among posterior regions rapidly decreased, and highly correlated networks among anterior motor areas (orange and red edges) subsequently emerged (**Figure 2B top**). The node degree of most areas rapidly declined after locomotion onset (15.4 ± 3.6 at 1 s, *n* = all 50 ROIs, mean ± SD), although the primary motor area remained elevated (M1p, 13.5 ± 0.7 at −3 s; 16.7 ± 0.7 at −1 s; 17.7 ± 0.1 at 1 s, *n* = 2 ROIs; **Figure 2C**). At the cessation of locomotion, the dense anterior networks among motor areas disappeared, and the FC among posterior regions reemerged (**Figure 2B middle**). These locomotion-dependent dynamic reconfigurations of functional network architecture were absent during random control periods (**Figure 2B bottom**). Taken together, the results demonstrate that the correlation among anterior motor areas becomes dominant over posterior sensory/association cortices during locomotion, whereas this reciprocal relationship between anteroposterior cortical domains is reversed during rest.

The betweenness centrality of M1p remained high during locomotion (**Figure 2D**), consistent with the notion that the primary motor area plays a pivotal role in voluntary movement. In addition, the betweenness centrality of the CG and PTa increased rapidly at locomotion onset, and PTa was high again immediately before cessation of locomotion. In contrast, TR and Vma displayed delayed rises after the onset and peaked immediately before cessation (**Figure 2D**). Fluorescence changes of each ROI were significantly correlated with node degree but not with betweenness centrality before locomotion onset (**Figure S4**), indicating that these functional network properties do not directly reflect the magnitude of fluorescence changes. These findings suggest that locomotion onset and cessation do not necessarily mirror each other, and hub structure dynamically changes within the period of locomotion. Moreover, we found significant increases in characteristic path length (CPL), a measure of the efficiency of information transfer that represents the average shortest path length between all region pairs (**Figure 2E**), and modularity Q, which means the extent to which the network is subdivided into nonoverlapping groups of regions (**Figure 2F**). These findings suggest that functional cortical networks manifest a more modular structure during movement than during rest periods of no locomotion. We found that correction for hemodynamic signals did not significantly impact these findings (**Figure S5**).

### Role of visual feedback in behavioral state-dependent cortical network dynamics

Animals use visual information to explore external environments. To investigate the role of visual sensory processing on our results, we tested mice exploring a virtual environment with no projection of visual landscape (**Figure 3**). In this condition, mice spent 46.9 ± 24.8 % and 35.2 ± 24.6 % of total time engaged in long locomotion and rest, respectively (mean ± SD; long locomotion, *P* = 0.38, vs. control; long rest, *P* = 0.46, vs. control, t-test, *n* = 71 sessions from 17 mice). Total distances traveled did not differ significantly from the control experiments with projection (**Figure 3B**). However, the mice often exhibited local circling and were impeded by invisible walls and corners when they explored without visual feedback (**Figure 3A**). As a result, they traversed a significantly smaller area within the arena (**Figure 3C**). These results demonstrate that vision provides important sensory information when mice explore the virtual arena.

**Figure 3.**
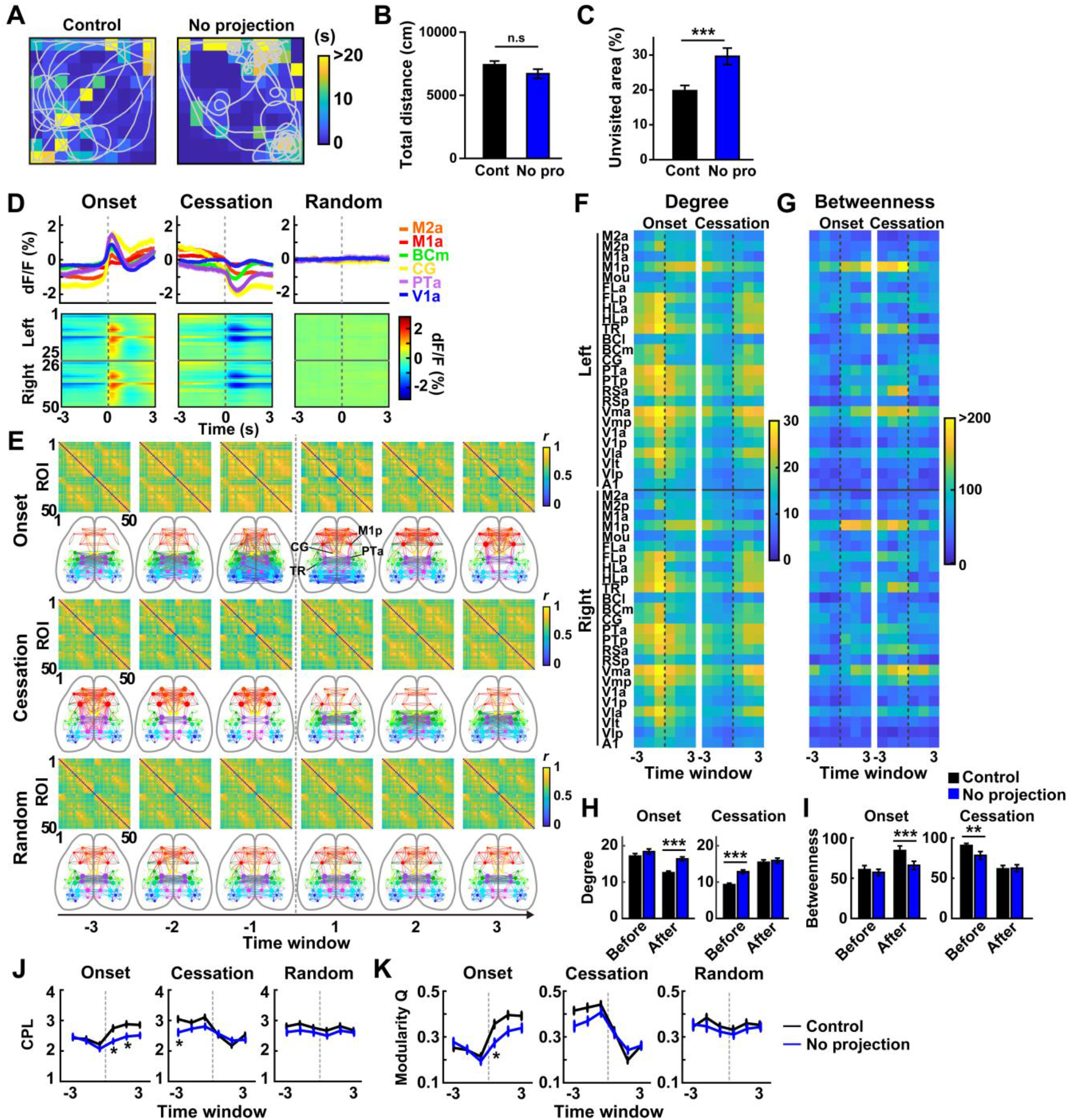
Changes in behavior and functional cortical network during exploration without visual feedback. (**A**) Representative trajectories overlaid onto heatmaps of dwell time during exploration with (Control) and without (No projection) visual feedback. (**B, C**) Distance traveled (B) and percentage of unvisited areas (C) during 10-min sessions with (Cont) or without (No pro) visual feedback. Data represent mean ± SEM. n.s., *P* = 0.14, \*\*\**P* < 0.001, t-test, *n* = 89 control sessions from 7 mice and 71 no projection sessions from 17 mice. **(C)** Cortical activity of Emx1G6 mice without visual feedback. The convention of the figure is the same as in Figure 2A. **(D)** Dynamics of correlations between activities of ROI pairs during the behavioral transitions without visual feedback. FC graphs (*r* > 0.8) were generated using the data shown in (D). The convention of the figure is the same as in Figure 2B. (**F, G**) Changes in node degree (F) and betweenness centrality (G) during the transitions in each ROI. (**H, I**) Mean node degree (H) and mean betweenness centrality (I) during the transitions. Data across all ROIs were averaged. \*\**P* < 0.01, \*\*\**P* < 0.001, t-test, *n* = 89 control sessions from 7 mice and 71 no projection sessions from 17 mice. (**J, K**) Change in CPL (J) and modularity Q (K) during the transitions. The control data presented in Figure 2 are again shown in black for comparison. Data represent mean ± SEM. (CPL) Onset, Time: *F*_(5, 930)_ = 11.95, P = 3.1×10^−11^; Genotype: *F*_(1, 930)_ = 22.28, *P* = 2.7×10^−6^; Time×Genotype: *F*_(5, 930)_ = 2.68, *P* = 0.02. Cessation, Time: *F*_(5, 924)_ = 19.59, *P* = 1.4×10^−18^; Genotype: *F*_(1, 924)_ = 9.47, *P* = 0.002; Time×Genotype: *F*_(5, 924)_ = 3.29, *P* = 0.006; Random, Time: *F*_(5, 948)_ = 1.25, *P* = 0.28; Genotype: *F*_(1, 948)_ = 9.65, *P* = 0.002; Time×Genotype: *F*_(5, 948)_ = 0.12, *P* = 0.99; (modularity Q) Onset, Time: *F*_(5, 930)_ = 29.78, *P* = 4.0×10^−28^; Genotype: *F*_(1, 930)_ = 11.03, *P* = 9.3×10^−4^; Time×Genotype: *F*_(5, 930)_ = 3.19, *P* = 0.007; Cessation, Time: *F*_(5, 924)_ = 44.90, *P* = 1.5×10^−41^; Genotype: *F*_(1, 924)_ = 4.37, *P* = 0.04; Time×Genotype: *F*_(5, 924)_ = 2.94, *P* = 0.01; Random, Time: *F*_(5, 948)_ = 1.45, *P* = 0.20; Genotype: *F*_(1, 948)_ = 2.78, *P* = 0.10; Time×Genotype: *F*_(5, 948)_ = 0.42, *P* = 0.83, two-way ANOVA. \**P* < 0.05, vs. control, Tukey Kramer test. *n* = 89 control sessions from 7 mice and 71 no projection sessions from 17 mice.

We then examined FC dynamics under no visual feedback. The fluorescence changes at locomotion onset and cessation were comparable to those in the control experiments except for larger and smaller amplitudes in CG (mean ± SD; 136.8 ± 34.0 %, vs. control, *n* = 71 sessions from 17 mice) and V1a (67.1 ± 19.9 %), respectively (**Figures 3D and 2A**). While FC networks were similar to those in the control experiments (**Figures 3E and 2B**), the number of connections during locomotion was significantly higher under no projection (**Figures 3F, 3H, and 2C**, after onset and before cessation). As in control experiments, the betweenness centrality of M1p was constantly high during locomotion. However, these centralities of CG, TR, and Vma were substantially reduced (no projection, 102.8 ± 41.7; control, 145.5 ± 33.7; mean ± SD, *n* = 6 ROIs; **Figures 3G and 2D**), which led to significant decreases in overall betweenness centrality during locomotion (**Figure 3I**). Furthermore, CPL and modularity Q were also significantly reduced during locomotion onset (**Figure 3J–3K**). These results indicate that the lack of visual feedback markedly weakens the network modularity of locomotion-dependent cortical FCs.

Desynchronization of cortical population activity often manifests when animals engage in a task ^22^. Using the network-based statistic (NBS) ^23^, we next tested for changes in FC during transitions between rest and locomotion. Importantly, this analysis allowed us to identify FCs that exhibited not only significantly higher correlations but also significantly lower correlations (i.e., decorrelation). In the control experiments, we observed gradual and marked increases in FC among posterior sensory areas and emergence of decorrelated subnetworks of anterior motor areas before locomotion onset (time window from –3 to –1, **Figure 4A**). After locomotion began, long-range decorrelations among anterior (motor areas), parietal (CG) and posterior (visual areas) cortices rapidly emerged, followed by robust decorrelations among posterior sensory cortices 2–3 s after locomotion onset (**Figure 4A**). In contrast, persistently decorrelated subnetworks of posterior sensory cortices during locomotion expanded to include M2 immediately before locomotion cessation, followed by the emergence of sustained decorrelations among anterior motor cortices and widespread transient correlations among posterior sensory cortices beginning 1–2 s after cessation (**Figure 4B**).

**Figure 4.**
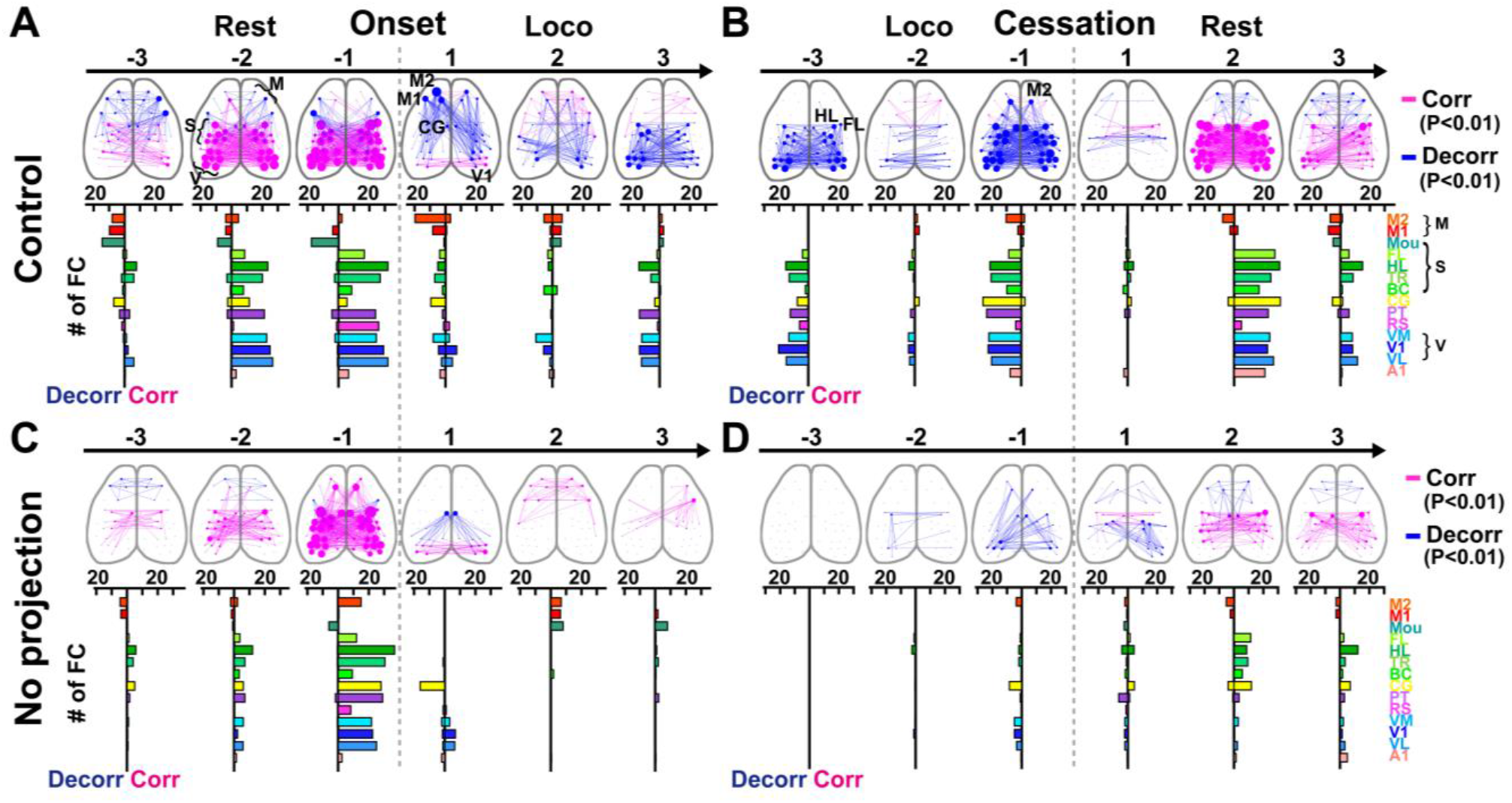
Statistically significant correlations and decorrelations within functional cortical networks during behavioral transitions with or without visual feedback. (**A, B**) Significant correlations and decorrelations of functional cortical subnetworks of Emx1G6 mice during locomotion onset in control experiments (A). Network diagrams of statistically significant FC during each second before and after the locomotion onset are shown from left to right (top). Magenta and blue lines denote significant correlations (Corr) and decorrelations (Decor) compared to the random control, respectively. The horizontal bar plots (bottom) indicate the number of significant FC (rightward: correlated, leftward: decorrelated) connected to each cortical area. The cortical areas are sorted along the antero-posterior axis from top to bottom. The values were averaged across bilateral ROIs and further averaged across multiple ROIs if the area contained more than one ROI. The same convention applies to locomotion cessation (B). Loco, locomotion; M, motor areas; S, somatosensory areas; V, visual areas. *P* < 0.01, NBS. (**C, D)** Significant correlations and decorrelations of functional cortical subnetworks during locomotion onset (C) and cessation (D) in Emx1G6 mice under no projection of visual landscape. *P* < 0.01, NBS.

The emergence of dense (de)correlated networks among sensory areas during behavioral transitions suggests that sensory processing could profoundly affect FC during these periods. Interestingly, we found that the decorrelated but not correlated networks markedly diminished in the condition with no projection of visual landscape (**Figures 4C and 4D**). Rapid decorrelations between M2 and V1 at ∼1 s after locomotion onset and delayed decorrelations among posterior sensory areas, including not only visual but also somatosensory cortices, at ∼3 s after locomotion onset were almost absent (**Figure 4C**), although decorrelations between CG and V1 and correlations between bilateral visual areas at ∼1 s after locomotion onset remained. Persistent decorrelations among posterior sensory areas before cessation and transient correlations among those areas ∼2 s after the cessation were considerably weakened (**Figure 4D**). Collectively, these results demonstrate that the absence of visual feedback markedly alters exploration behavior and dynamics of multiple functional subnetworks primarily involving the visual cortex, such as long-range anteroposterior FCs between motor and visual cortices and cross-modal FCs between somatosensory and visual cortices.

### Decoding behavioral dynamics using functional cortical network

Having found that the transitions between states of rest and locomotion were each characterized by distinct cortical network architectures, we next tested whether an animal’s behavioral state can be decoded from its cortical network. To this end, we trained support vector machine (SVM) classifiers using datasets of FC containing all time frames from four mice (train set) and classified FCs for the remaining three mice (test set) into two behavioral states. This was repeated for all combinations of mouse assignments to test and train sets. Accuracy of the out-of-sample classification (Test, 88.9%, median, *n* = 35 classifiers) was comparable to the level achieved by classification for the train set (Train, 89.0%) and substantially higher than expected due to chance, as determined by randomly shuffling classification labels (Shuffled, 58.3%; **Figure 5A**). Incorrect classification mainly occurred during short periods flanking the state transition, and classification accuracy was highest during continuing locomotion and rest periods. However, accuracies remained modest during the intermediate periods in which two contrasting states coexisted (time window from −1 to 1, **Figure 5B**). Accordingly, the accuracy of classification increased to 92.3 % (*n* = 1,326,379 frames from 89 sessions) when the periods of short locomotion and rest episodes (less than 3 s) were excluded, whereas accuracy within the periods of short episodes was 65.0 % (*n* = 275,621 frames from 89 sessions).

**Figure 5.**
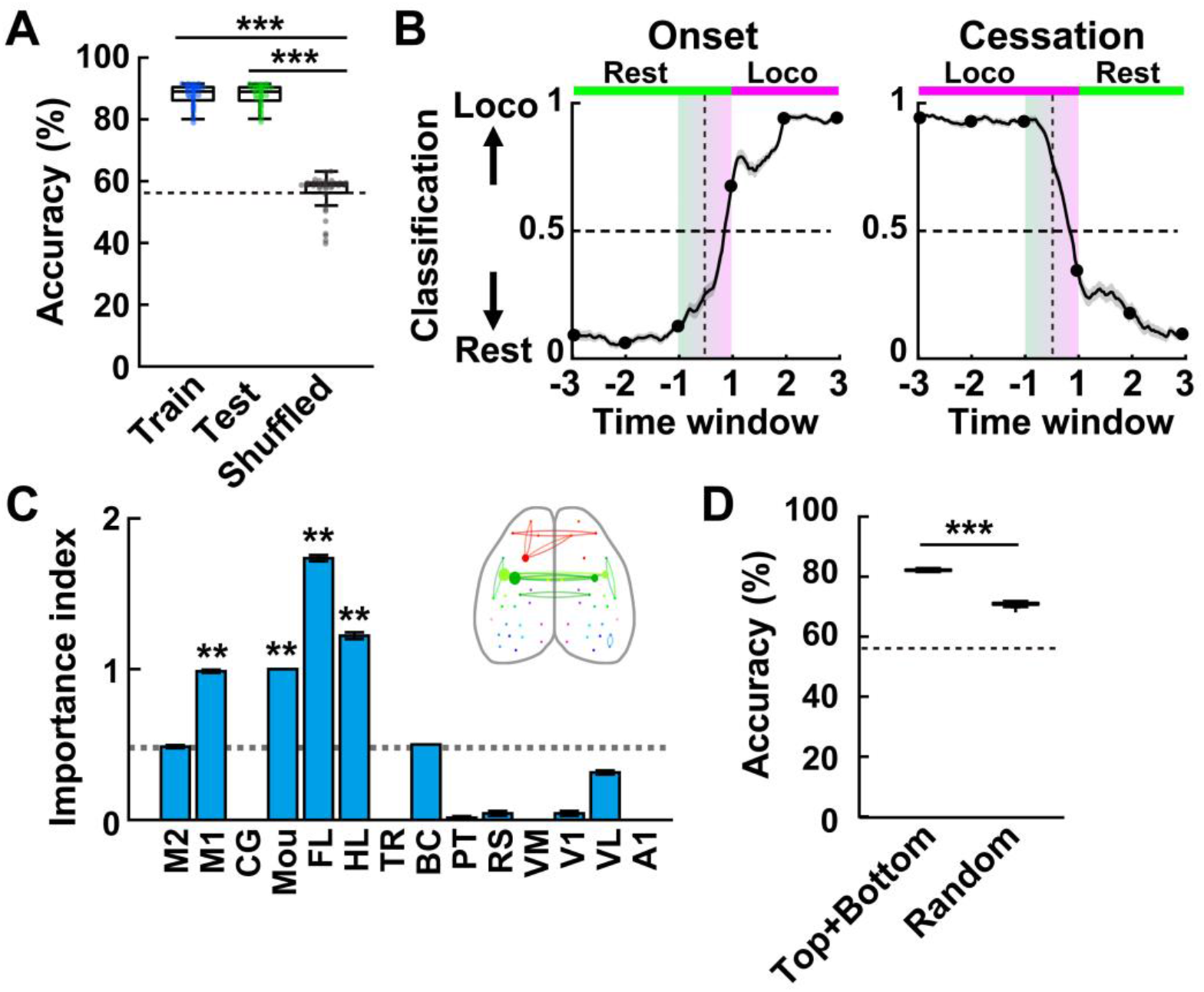
Decoding behavioral states from the functional cortical network on a subsecond time scale. **(A)** Accuracy of SVM classification of FC into the two behavioral states. The results of classification for train set (Train), test set (Test), and shuffled control (Shuffled) are shown. Data represent averages across entire sessions (17,970 time points each). The boxes represent the 25th, 50th, and 75th percentiles, and the whiskers represent the range except for outliers. The dashed line indicates the chance level defined as an overall average of time spent in locomotion (58.7 %). \*\*\**P* < 0.001, vs. Shuffled. Wilcoxon rank-sum test with Bonferroni correction. *n* = 35 classifiers each. **(B)** Dynamics of SVM classification of FC during locomotion onset and cessation. Y-axis indicates classification index (1, classified into locomotion state; 0, classified into resting state). The magenta and green bars on the top indicate the periods during locomotion (Loco) and rest, respectively. The gradation bar from -1 to 1 indicates that the time window during this period contained FC from both locomotion and rest periods. Data represents mean ± SEM (*n* = 35 classifiers). **(C)** Importance index of each cortical area. The index was defined as the number of the appearance of FCs connected to each area in the top 0.5 % and bottom 0.5 % features (see STAR Methods for details). The dashed line indicates a chance level defined as an average of 100-times random sampling of 1 % features. (Inset) Functional networks of the top 0.5 % and bottom 0.5 % features (important index: ≥ 0.2). \*\**P* < 0.01, vs. chance level. Wilcoxon rank-sum test with Bonferroni correction. *n* = 35 classifiers each. **(D)** Classification accuracy using the top 0.5 % and bottom 0.5 % features (Top+Bottom) and randomly selected 1 % features (Random). \*\*\**P* < 0.001, vs. Random. Wilcoxon rank-sum test. *n* = 35 classifiers each.

To identify the features that contributed significantly to the classification, we sorted all FCs according to feature weights and found that M1, Mou, FL, and HL were significantly overrepresented in the top 0.5 % and bottom 0.5 % FCs (6 FCs each) (**Figure 5C**). We then retrained the classifiers using these top 0.5 % and bottom 0.5 % FCs and achieved classification accuracies that were comparable to the classifier trained with all FCs (Top+Bottom, 84.2 %; **Figure 5D**) and significantly better than the classifiers trained with a randomly selected 1 % of all FCs (Random, 71.0 %; **Figure 5D**). Collectively, these results demonstrate that connectivity of the primary motor and primary somatosensory forelimb, hindlimb, and mouth areas contains information sufficient for highly accurate differentiation of locomotion and rest states.

### Functional hyperconnectivity and impaired locomotion-dependent dynamics in the cortex of a mouse model of ASD

We applied our VR-based imaging system to investigate behavior-dependent cortical network dynamics of ASD model mice. We used Emx1G6_*15q dup*_ mice that possessed the paternal duplication of the mouse syntenic region of human 15q11-13 and expressed GCaMP6f in excitatory neurons in the cortex. Emx1G6_*15q dup*_ mice showed lower locomotor activity in the virtual arena during 10-min sessions (**Figures 6A and 6B**). They spent 25.8 ± 17.1 % and 59.1 ± 21.3 % of total time engaged in long locomotion and rest, respectively (mean ± SD; long locomotion, *P* = 5.3×10^−14^, vs. Emx1G6; long rest *P* = 6.9×10^−15^, vs. Emx1G6; t-test, *n* = 88 sessions from 9 mice; **Figure 6C**), and average lengths of long locomotion and rest per episode were 9.0 ± 4.2 s (*P* = 0.17, vs. Emx1G6, t-test, *n* = 1,523 episodes from 88 sessions) and 20.5 ± 14.7 s (*P* = 2.7×10^−9^, vs. Emx1G6, t-test, *n* = 1,882 episodes from 88 sessions), respectively. Functional sensory mapping confirmed that locations and response amplitudes of primary somatosensory subareas were not markedly different between Emx1G6_*15q dup*_ mice and Emx1G6 mice (**Figure S7**), although the area responsive to whisker stimuli was larger in Emx1G6_*15q dup*_ mice (**Figures S7C and S7D**), as reported in our previous study ^20^.

**Figure 6.**
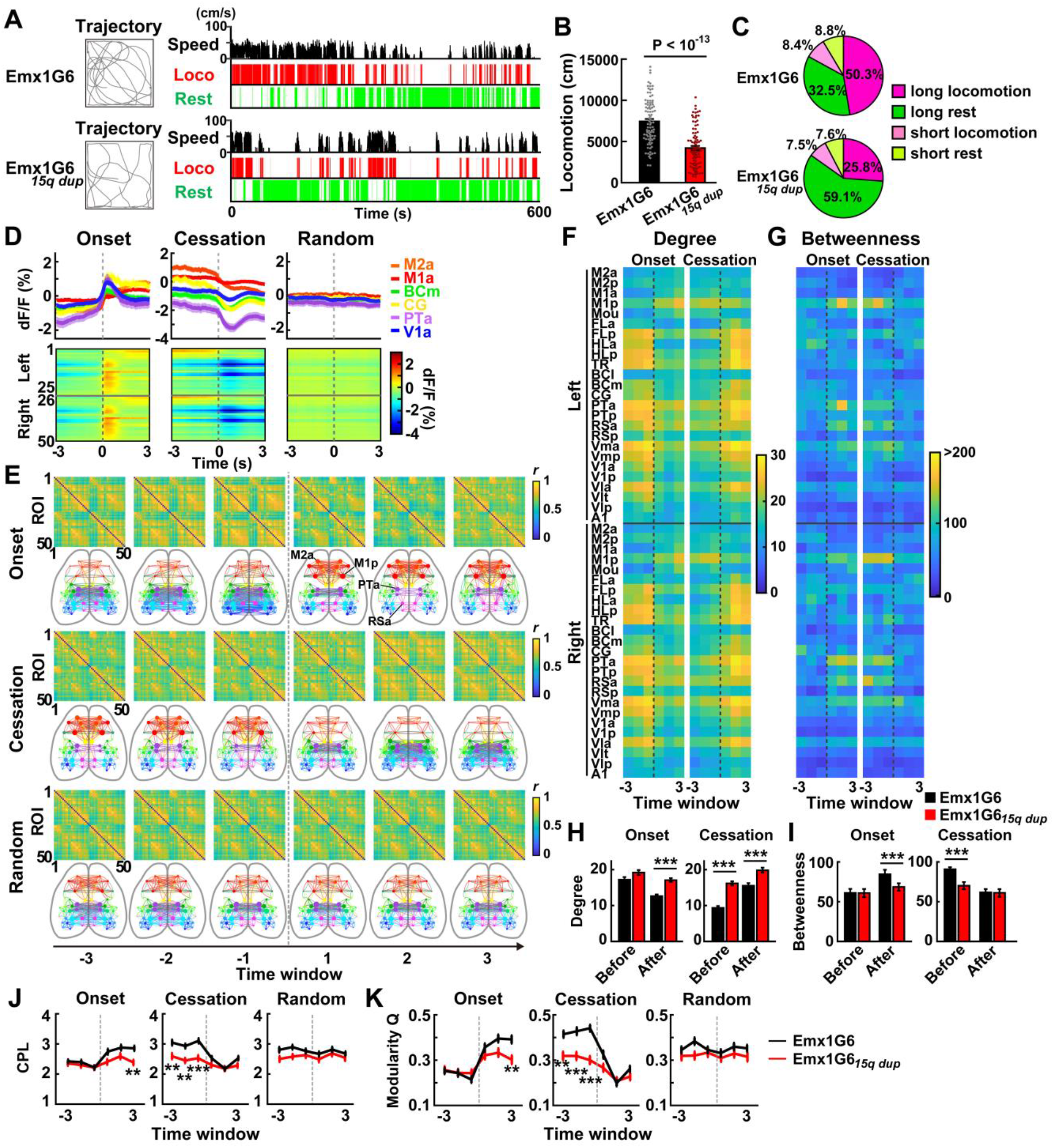
Hyperconnectivity and lower modularity of functional cortical networks in Emx1G6_*15q dup*_ mice during behavioral state transitions. **(A)** Representative trajectory (left) and locomotion behavior (right) for Emx1G6 mice and Emx1G6_*15q dup*_ mice. Locomotion speed, periods of locomotion (Loco), and the rest of each genotype are shown from top to bottom in the right panel. **(B)** Locomotor activity of Emx1G6 mice and Emx1G6_*15q dup*_ mice during 10-min sessions. Data represent mean ± SEM. *P*-value by t-test. *n* = 89 sessions from 7 Emx1G6 mice and 88 sessions from 9 Emx1G6_*15q dup*_ mice. **(C)** Percentages of time spent for each episode in Emx1G6 mice and Emx1G6_*15q dup*_ mice. Data represent averages across all sessions. **(D)** Cortical activity of Emx1G6_*15q dup*_ mice during the behavioral transitions. The convention of the figure is the same as in Figure 2A. **(E)** Dynamics of correlations between activities of ROI pairs during the transitions in Emx1G6_*15q dup*_ mice. FC graphs (*r* > 0.8) were generated using the data shown in (D). The convention of the figure is the same as in Figure 2B. (**F, G**) Changes in node degree (F) and betweenness centrality (G) during the transitions in each ROI of Emx1G6_*15q dup*_ mice. (**H, I**) Mean node degree (H) and mean betweenness centrality (I) during the transitions. Data across all ROIs were averaged. \*\*\**P* < 0.001, t-test, *n* = 89 sessions from 7 Emx1G6 mice and 88 sessions from 9 Emx1G6_*15q dup*_ mice. (**J, K**) Change in CPL (J) and modularity Q (K) during the transitions in Emx1G6_*15q dup*_ mice. The data for Emx1G6 mice presented in Figure 2 are again shown in black for comparison. Data represent mean ± SEM. (CPL) Onset, Time: *F*_(5, 1014)_ = 10.35, *P* = 1.1×10^−9^; Genotype: *F*_(1, 1014)_ = 19.09, *P* = 1.4×10^−5^; Time×Genotype: *F*_(5, 1014)_ = 2.30, *P* = 0.04. Cessation, Time: *F*_(5, 1026)_ = 19.60, *P* = 1.1×10^−18^; Genotype: *F*_(1, 1026)_ = 47.70, *P* = 8.7×10^−12^; Time×Genotype: *F*_(5, 1026)_ = 3.37, *P* = 0.005; Random, Time: *F*_(5, 1050)_ = 1.28, *P* = 0.27; Genotype: *F*_(1, 1050)_ = 16.18, *P* = 6.2×10^−5^; Time×Genotype: *F*_(5, 1050)_ = 0.49, *P* = 0.78; (modularity Q) Onset, Time: *F*_(5, 1014)_ = 25.25, *P* = 4.7×10^−24^; Genotype: *F*_(1, 1014)_ = 6.73, *P* = 0.01; Time×Genotype: *F*_(5, 1014)_ = 3.53, *P* = 0.004; Cessation, Time: *F*_(5, 1026)_ = 38.98, *P* = 1.0×10^−36^; Genotype: *F*_(1, 1026)_ = 53.33, *P* = 5.6×10^−13^; Time×Genotype: *F*_(5, 1026)_ = 5.49, *P* = 5.4×10^−5^; Random, Time: *F*_(5, 1050)_ = 0.82, *P* = 0.54; Genotype: *F*_(1, 1050)_ = 10.23, *P* = 0.001; Time×Genotype: *F*_(5, 1050)_ = 0.52, *P* = 0.76, two-way ANOVA. \*\**P* < 0.01, \*\*\**P* < 0.001, vs. Emx1G6, Tukey Kramer test. *n* = 89 sessions from 7 Emx1G6 mice and 88 sessions from 9 Emx1G6_*15q dup*_ mice.

Although the fluorescence changes at locomotion onset and cessation in Emx1G6_*15q dup*_ mice were generally similar to those in Emx1G6 mice (**Figure 6D**, see also Figure 2A), the magnitude of changes in a few areas, such as PTa (mean ± SD; Emx1G6_*15q dup*_, -0.10 ± 0.21 %; Emx1G6, 1.33 ± 0.29 % at 0 s) and BCm (Emx1G6_*15q dup*_, -0.15 ± 0.20 %; Emx1G6, 0.63 ± 0.15 % at 0 s), were low (**Figure 6D**). These differences were not likely due to different baseline fluorescence levels in Emx1G6_*15q dup*_ mice, as average fluorescence intensities of each cortical area were only slightly higher in these mice (**Figure S6**). The overall patterns of FC networks in Emx1G6_*15q dup*_ mice were also similar to those in Emx1G6 mice (**Figure 6E**, see also Figure 2B). However, the strength of FC appeared higher in Emx1G6_*15q dup*_ mice, particularly FCs connecting ROIs in anterolateral motor cortices (i.e., M1 and M2) and FCs bridging anterior and posterior cortex (e.g., M1, PT, and RS) during locomotion (**Figure 6E**), as supported by generally larger node degrees compared with Emx1G6 mice (**Figures 6F and 6H**). In Emx1G6_*15q dup*_ mice, betweenness centrality, CPL, and modularity Q were significantly lower than in Emx1G6 mice during locomotion (**Figures 6I–6K**), suggesting that cortical hyperconnectivity results in a less modularized, more interconnected network in behaving Emx1G6_*15q dup*_ mice. Given the baseline hyperconnectivity in Emx1G6_*15q dup*_ mice, surprisingly fewer locomotion-dependent decorrelations were detected, particularly in the posterior cortex (**Figure S8**). These findings collectively demonstrate that functional cortical networks in Emx1G6_*15q dup*_ mice exhibit hyperconnectivity and enhanced interconnectivity, but a locomotion-related reconfiguration of network architecture is dampened compared to Emx1G6 mice.

### Diagnosis of ASD model mice using temporal FC during behavioral transitions

Compared to Emx1G6 mice, we found that Emx1G6_*15q dup*_ mice were characterized by significant hyperconnectivity of M2 and M1 areas during the transitions, most notably 1 s after onset and 2 s after cessation of movement (**Figures 7A and 7B**). Bilateral connections of somatosensory nodes (especially HL and TR) were significantly decorrelated compared with Emx1G6 mice regardless of behavioral state (**Figures 7A and 7B**). In addition, FCs between PT and M1, M2, FL, and HL showed decorrelation during locomotion, most evidently before cessation (**Figure 7B**).

**Figure 7.**
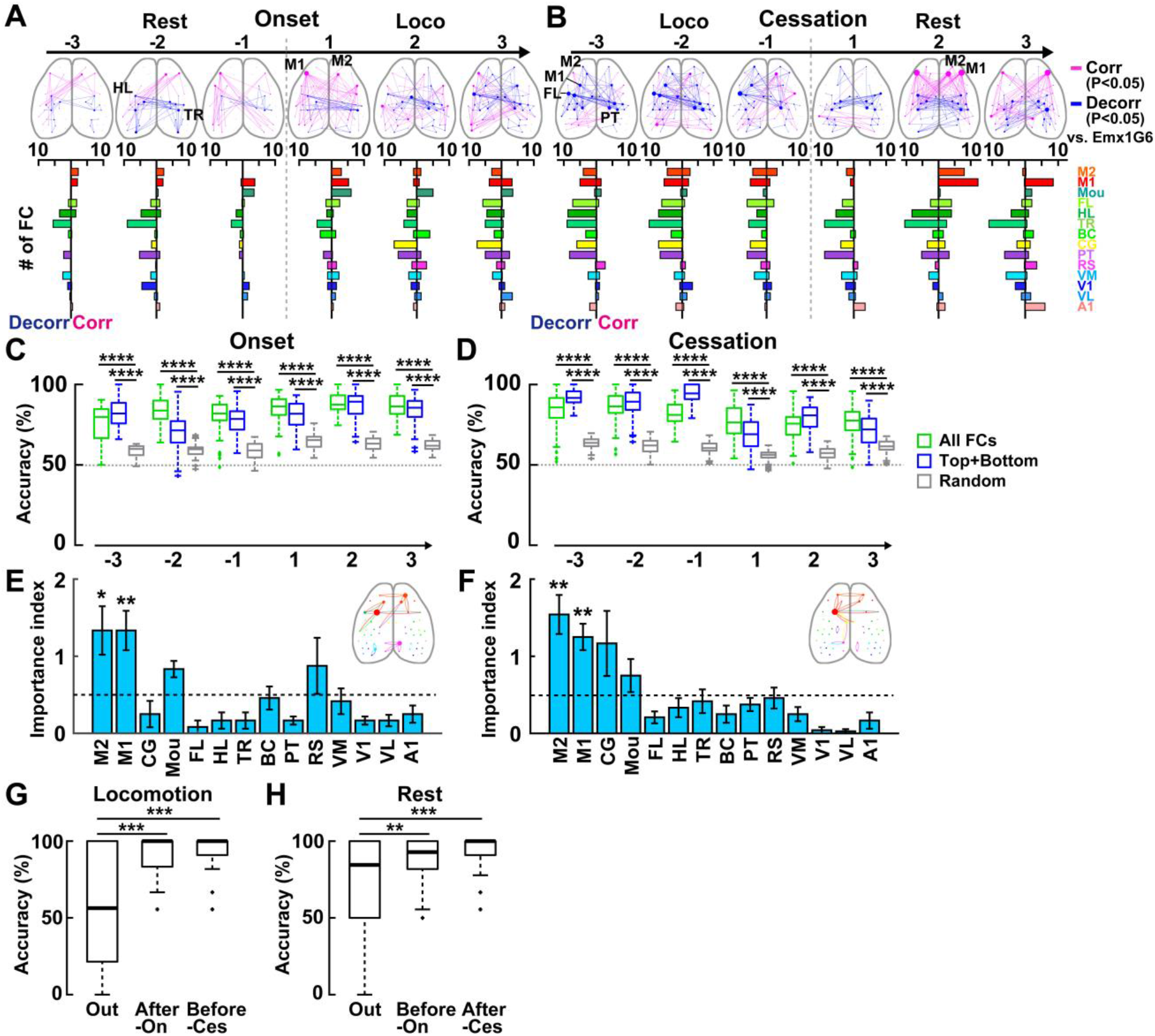
Importance of motor areas and behavioral transitions in distinguishing cortical FC between Emx1G6_*15q dup*_ and Emx1G6 mice. (**A, B**) Statistically significant FC of Emx1G6_*15q dup*_ mice during locomotion onset (A) and cessation (B) compared to Emx1G6 mice. The convention of the figure is the same as in Figure 4A. *P* < 0.05, NBS. (**C, D**) Accuracy of SVM classification of FC into two genotypes during locomotion onset (C) and cessation (D). Classifiers were trained with all 1,225 features (All FCs), top 0.5 % and bottom 0.5 % features (Top+Bottom), or randomly chosen 1 % features (Random) at each time point. The boxes represent the 25th, 50th, and 75th percentiles, and the whiskers represent the range except for outliers. \*\*\*\**P* < 0.001, vs. Random, Wilcoxon rank-sum test with Bonferroni correction. *n* = 63 classifiers each. (**E, F**) The importance index of each cortical area in the SVM classification of FC during locomotion onset (E) and cessation (F) was averaged across all relevant time points. Data represent mean ± SEM (*n* = 6 time points). The dashed line indicates a chance level defined as an average of 100-times random sampling of 1 % features. (Inset) Functional networks of the top 0.5 % and bottom 0.5 % features (≥ 2 time points). \**P* < 0.05, \*\**P* < 0.01, vs. chance level, Wilcoxon rank-sum test with Bonferroni correction. (**G, H**) Accuracy of SVM genotype classifiers trained with FC during locomotion (G) and rest (H) with or without transitions. The periods of locomotion were subdivided into those that occurred immediately after locomotion onset (After-On), immediately before locomotion cessation (Before-Ces), and outside these two types of periods (Out). Similarly, the periods of rest were subdivided into those that occurred immediately before locomotion onset (Before-On), immediately after locomotion cessation (After-Ces), and outside these two types of periods (Out). \*\**P* < 0.01, \*\*\**P* < 0.001, vs. Out, Wilcoxon rank-sum test with Bonferroni correction. *n* = 63 classifiers each.

To identify FC features that most distinguished Emx1G6_*15q dup*_ mice from Emx1G6 mice, we conducted SVM classification of FC during locomotion onset and cessation into the two genotypes. The SVM classifiers trained with all features (All FCs) at each time point during behavior transitions accurately classified the Emx1G6_*15q dup*_ and Emx1G6 genotypes (**Figures 7C–7F**). As with behavior state classification, the SVM classifiers trained with top 0.5 % and bottom 0.5 % features (Top+Bottom) performed comparably to the classifiers trained with all FCs, and significantly more accurately classified FC than the classifiers trained with randomly selected 1 % features (Random) at all time points (**Figures 7C and 7D**). FCs including M2 and M1 were significantly over-represented in the top 0.5% and bottom 0.5% features (**Figures 7E and 7F**), pointing to these cortical areas as key nodes that primarily contribute to deficits of cortical processing during spontaneous behavioral switching of Emx1G6_*15q dup*_ mice.

Finally, we tested the importance of the behavior transition periods for genotype classification. The accuracies of classifiers trained with data from the locomotion that occurred within the transition periods (median accuracy, After-On, 100 %; Before-Ces, 100 %; **Figure 7G**) were significantly higher than those trained with data from continuous locomotion outside the transition (Out, 56.3 %; **Figure 7G**). Similarly, classifiers trained with datasets from the rest that occurred within the transition periods more accurately classified FC into the right genotype than classifiers trained with the data from continuous rest periods outside the transition (Before-On, 92.9 %; After-Ces, 100 %; Out, 84.5 %; **Figure 7H**). In summary, these results demonstrate that the distinguishability of FC is greater during transition periods than during continuous locomotion and rest.

## Discussion

In this study, we investigated locomotion-induced changes in rapid cortico-cortical FC on the time scale of seconds by taking advantage of an integrated platform for mesoscopic Ca^2+^ imaging and VR that allows mice to run spontaneously with sensory feedback. Neural activity signals obtained using fluorescent Ca^2+^ indicator proteins are faster and typically more spatially resolved than BOLD signals. FC measured using fMRI and mesoscopic functional imaging is shown to overlap mainly with underlying structural connectivity ^24–26^ and reflect the correlation of modulation of neuronal spiking and LFP (local field potential) power between brain regions ^27–29^. Our correlation-based FC analysis thus highlighted communication and interaction between cortical areas based on the level of local activity.

### Cortical FC dynamics during behavioral transitions with and without visual feedback

The locomotion-dependent cortical functional network changes revealed in this study align with previous observations ^27,30–34^. Recent imaging studies demonstrate that M2, which has dense reciprocal anatomic connections with sensory, parietal, and retrosplenial cortices ^35^, orchestrates widespread cortical activity during motor learning and in a decision-making task ^30,33^. In our study, in line with the view that M2 acts to link antecedent conditions such as sensory information to motor actions ^36^, M2 showed a transient elevation of significant correlation with multiple sensory areas at 1–2 s ahead of locomotion onset, regardless of the presence or absence of visual feedback (Figures 4A and 4C). Recently, a study that investigated cortical FC dynamics during locomotion also highlighted the importance of M2 ^34^. However, the role of sensory feedback for FC in this node had not been directly examined. Here, we found that FC between M2 and sensory cortices, including primary somatosensory cortex (S1) and primary visual cortex (V1), was decorrelated at 1 s after locomotion onset and demonstrated that this decorrelation completely disappeared when visual feedback was not available (Figures 4A and 4C). The implication is that locomotion with visual feedback drives V1 more strongly than without feedback and that direct top-down input from M2 to V1 sends motor-related signals for visual flow predictions ^31,32^.

The FC associated with S1 significantly contributed to the SVM classification of locomotion and rest (Figure 5C). Remarkably, a dense correlated network among nodes of sensory areas, including S1, exhibited widespread and gradual augmentation over a period of 2 s before locomotion onset, but this characteristic functional subnetwork was no longer evident once locomotion started (Figure 4A). This preparatory emergence of a correlated network is reminiscent of the synchronous oscillations observed in S1 during premovement attentive immobility ^37,38^ and is also consistent with the recent finding that S1 neuronal activity is highly correlated with the onset of movement and can control locomotion through a direct pathway independently of the motor cortex ^39^. Our analysis of fast FC dynamics was thus able to capture a global picture of distributed transient functional subnetworks that may play a role in the preparation and initiation of voluntary movement.

### Cortical FC abnormalities in *15q dup* mice

Our FC analysis of a mouse model of ASD uncovered previously unknown impairment of cortical circuit function such as widespread hyperconnectivity, less modularized network during locomotion, and FC patterns involving M2 and M1 as the most distinctive signature for *15q dup* mice.

It has been reported that individuals with ASD exhibit motor coordination deficits and impairment of movement planning in goal-directed locomotion ^10–12^. Although various factors could influence the locomotor activity of mice, reduced time spent for long locomotion in Emx1G6_*15q dup*_ mice might result from impaired motor planning and execution due to abnormal M2-related FC. While human dup15q syndrome shows a gait pattern of the slow pace, poor postural control, and large gait variability ^40^ and patients with paternal duplication in 15q11-13 display clumsy motor skill development ^41^, *15q dup* mice were also reported to display mild motor impairment such as longer stride length and reduced stride frequency, and deficits in motor learning and cerebellar synaptic plasticity ^42^. Since recent studies demonstrate that cerebellar output modulates preparatory activity in the anterolateral motor cortex ^43,44^, the abnormal M2-related FC we observed during behavior state transitions may also arise as a consequence of deficiency of a more widespread functional network, potentially including interactions with extracortical brain regions.

Compared with Emx1G6 mice, Emx1G6_*15q dup*_ mice show significant decorrelation of FC that links M2, CG, S1, and PT during locomotion (Figures 7A and 7B). This subnetwork is reminiscent of the human lateral frontoparietal network (L-FPN), which consists of the rostral and dorsolateral prefrontal cortex and the inferior parietal cortex and participates in executive functions such as goal-directed cognition and task switching ^45^. In task-based fMRI studies, atypical activation of L-FPN is observed during cognitive flexibility tasks in ASD brains ^46^. Thus, it would be of interest in the future to investigate whether abnormal interaction between nodes of a mouse L-FPN equivalent in *15 dup* mice is implicated in impaired behavioral flexibility observed in reversed learning tests of the Morris water maze and Barnes maze ^18^.

### Future outlook

Our machine learning classification results demonstrate that information regarding an animal’s ongoing behavioral state is represented in the fast dynamics of global cortical FC patterns (Figure 5). Identification of brain activity-based ASD biomarkers and machine learning-assisted diagnosis of ASD using neuroimaging data are fields of active investigation ^8,9,47^. While recent human fMRI studies have begun to explore the use of dynamic resting-state FC to identify atypical brain network activity unique to ASD ^46,47^, our results highlight the importance of examining behavioral transitions rather than simply looking at the resting state (Figures 7C–7F). Exploring additional mouse models will accumulate more evidence to identify common FC changes beyond heterogeneity of ASD ^48^. Furthermore, in future studies, it is of great interest to investigate whether the observed FC abnormalities can be reversed by pharmacologic treatment during postnatal development or adulthood of ASD model mice. Thus, our system to examine locomotion-dependent rapid FC changes based on mesoscopic cortex-wide Ca^2+^ imaging and VR offers a new translational approach toward developing precise diagnostic tools and effective treatment for various brain disorders. A fascinating future possibility would be to create a multimodal “metaverse” in which mice interact with other conspecifics via their avatars to understand cortical FC dynamics during virtual social interaction ^49^.

## STAR Methods

### Mice

The following Cre driver, reporter, and ASD model mouse lines were used for breeding; Emx1cre (B6.129P2-Emx1<tm1.1(cre)Ito>/ItoRbrc, RBRC01345, RIKEN Bioresource Center; ^50^), Ai95D (B6;129S-Gt(ROSA)26Sortm95.1(CAG-GCaMP6f)Hze/J, JAX024105, Jackson Laboratories), *15q dup* (B6.129S7-Dp(7Herc2-Mkrn3)1Taku, RBRC05954, RIKEN Bioresource Center; ^18^). Although some genotypes of transgenic mice that express GCaMP6 reportedly exhibit cortical epileptiform fluorescence events (most often seen in Ai93 line) ^51^, we did not observe such aberrant activity in our combination of Emx1-cre mice and Ai95D mice. For experiments, Emx1G6 mice were obtained by crossing Emx1-cre mice with Ai95D mice. Emx1G6_*15q dup*_ mice were obtained by crossing male mice double-positive for Emx1-Cre and *15q dup* and female mice positive for Ai95D. All mice were maintained in a reverse 12 h dark/light cycle (light off at 8 a.m.), and experiments were conducted during the dark phase. Food and water were available *ad libitum*.

### Surgery

All procedures were carried out following the institutional guidelines and protocols approved by the RIKEN Animal Experiments Committee. Twenty-three Emx1G6 mice, nine Emx1G6_*15q dup*_ mice, and three C57BL/6J (non-G6) mice (all male at 12–20 weeks old) were used for experiments. During surgery, mice were anesthetized under 1.5–2.0 % isoflurane in air, and the body temperature was kept at 37°C using a heating pad. The scalp was cut off and the surface of the skull was cleaned using a cotton swab. The skull surface was then covered with a thin layer of transparent resin (Super-Bond C&B, Sun Medical, Japan), followed by placement of a coverslip (0.17 mm thickness, Matsunami, Japan) onto the resin layer ^52^. A custom-made metal head plate with a polygonal imaging window (size of opening, 13 mm long and 10 mm wide, Narishige, Japan) was affixed to the edge of the coverslip with dental cement so that the entire dorsal cortex was clearly visible transcranially through the window (Figures 1C and 1G). The mice were allowed to fully recover from anesthesia in a warmed chamber and then returned to their home cages.

### VR environment

A VR environment for head-fixed mice was constructed as previously described with modification (Figures 1A and 1B) ^16,17^. An air-floated spherical treadmill was composed of a 20-cm polystyrene foam ball and a hemispherical stainless steel bowl with an internal diameter that fitted with the ball. The bowl had eight holes for pressured air at the bottom. The head of the mouse was fixed to a rigid head mount bar and posts via the head plate and positioned ∼1 cm above the top of the ball. The movement of mice was detected as rotations of the treadmill by two USB optical motion detectors (Gaming Mouse G302, Logicool) which were positioned orthogonal to each other on the equator of the treadmill. The movement signals from the motion detectors were transformed into analog output voltages using a custom-written LabVIEW program (National Instruments) to control the mouse’s virtual position via a joystick controller (USB Joystick Interface, 909991, APEM) connected to the VR software (OmegaSpace ver 3.7, Solidray). An interactive VR landscape rendered from a first-person perspective was projected by two compact liquid crystal display projectors (M110, Dell) onto the back of a custom-made 40 cm-diameter translucent acrylic semi-domal screen that was positioned 20 cm in front of the mouse and covered 240° of the mouse’s visual field.

### Behavioral testing and mesoscopic cortical-wide Ca^2+^ imaging

Mice underwent three pre-training steps to acclimate to the test environment. In the first step that began 3–5 days before surgery, mice were daily allowed to move freely on the top of the polystyrene foam ball that was rotated manually by an experimenter for 10 min and then handled by the experimenter under room light for another 10 min. The second step started as early as a day after surgery. In this step, mice were acclimated daily to head-fixation in the VR set-up for 3–5 days until they were able to sit and move on the treadmill in a balanced manner for 0.5–1 h under dim light (approximately 20 lux). In the final step, mice were acclimated to a complete VR environment and allowed to explore the virtual arena for 10 min daily for 5–10 days until they could move along the wall of the arena and turn the corner without difficulty.

After completing these pre-training processes, spontaneous locomotion within the virtual arena and cortical activity were recorded in a 10-min test session daily for a total of 15 sessions. The cortex was illuminated transcranially by a mercury lamp (U-HGLGPS, Olympus) through 460-480 nm (MGFPHQ, Olympus) or 457-487 nm (GFP-3035D, Semrock) excitation filters. Green fluorescence images were acquired using a CMOS camera (ORCA-Flash 4.0 v2, Hamamatsu) mounted on a HyperScope upright microscope (Scientifica) through a 2× objective lens (Plan Apo λ, NA: 0.10, Nikon) and 495-540 nm (Olympus) or 502.5-537.5 nm (Semrock) emission filters. Images of 512 × 512 pixels (14.8 μm × 14.8 μm/pixel; field of view, 7.5 mm × 7.5 mm) were collected at a rate of 30 frames per second while the head-fixed mouse freely explored the virtual arena. The mouse’s locomotion speed and coordinates were recorded at a sampling rate of 60 Hz using custom LabVIEW software. The rising edge of the TTL (Transistor-transistor-logic) signals that the camera generated at the acquisition of each frame were detected and recorded simultaneously with the behavioral data for synchronization with the imaging data. Experiments without projection of VR landscape were conducted (5 sessions after final sessions in the normal condition) by turning off the LCD projectors.

### ROI selection

A total of 50 ROIs were defined bilaterally (25 ROIs for each hemisphere) so that they covered all the cortical subregions designated in a dorsal cortical map ^53,54^ (Figure 1G). During our preliminary analysis, we visually inspected sample fluorescence movies of spontaneous cortical activity from three mice and selected several tens of ROI candidates that appeared brighter or darker than their surrounding regions. We then carefully examined and modified them so that the entire ROI set accords well with known cortical parcellations provided by annotated brain atlases ^53,54^. The resultant ROI map was registered with fluorescence images of the dorsal cortex by manual translation and rotation so that Bregma and the midline of the ROI map and fluorescence images were in the register. Each ROI was defined as a square of 5 × 5 pixels (within 128 × 128 pixel images) to avoid potential signal contamination across areal borders. In some cases, multiple ROIs assigned to relatively large cortical areas (e.g., primary somatosensory cortex, visual cortex, etc.) were arranged so that each corresponded to anatomical/functional subdivisions designated in the brain atlases.

The validity of our ROI positions for the primary somatosensory and primary visual cortices was confirmed by mapping sensory responses (Figure S1A). An air-puff (20 psi, 200 ms duration, PLI-10, Warner Instruments) to the right whiskers, forelimb, hindlimb, or the right side of the trunk and a flash of a yellow LED (0.2 Hz, 5 ms duration, Spectralynx, Neuralynx) to the right eye were given to mice anesthetized with 1.0–1.2 % isoflurane as tactile and visual stimuli, respectively, and the areas that displayed the largest average fluorescence changes calculated from 25–30 responses were compared to the corresponding ROIs. The validity of ROI positions for motor areas was confirmed by constructing a pixel-based correlation map between fluorescence changes of the pixel and locomotor activity (Figures S1B and S1C). The consistency of ROI registration processes within and across genotypes was validated by consistent positions of multiple ROIs that corresponded to the primary somatosensory subareas (*n* = 9–11 Emx1G6 mice and 4–5 Emx1G6_*15q dup*_ mice; Figure S7).

### Data analysis

For locomotion analysis, the locomotion speed recorded at 60 Hz was downsampled to 30 Hz to match the timing of image acquisition. Periods of locomotion were defined as those during which the locomotion speed exceeded 0.5 cm/s, and the other periods were defined as those of rest. Episodes of locomotion and rest that were equal to or longer than 3 s were then labeled as “long locomotion” and “long rest”, respectively. The remaining episodes were categorized as “short locomotion” and “short rest”. The threshold of 3 s was close to the average lengths of all locomotion and rest episodes (locomotion, 3.7 ± 7.3 s; rest, 2.6 ± 6.4 s; mean ± SD, *n* = 89 sessions) and was chosen to obtain a sufficient number of transition events per session (locomotion onset, 6.4 ± 3.9 events/session; locomotion cessation, 7.3 ± 4.7 events/session; mean ± SD, *n* = 89 sessions) while excluding periods of frequent alterations of the behavioral state that were too short to be used for the subsequent analysis of functional connectivity (FC) (Figure 1E). The exclusion of these periods did not likely affect the comparisons between Emx1G6 mice and Emx1G6_*15q dup*_ mice, as stereotypy measured in an open-field test ^55^, and the percentages of time spent on short locomotion and short rest were comparable between these genotypes (Figure 6C).

Raw fluorescence movies were spatially binned to 128 × 128 pixels and registered manually using ImageJ (NIH) so that the cortical image was aligned to a representative overhead view of the dorsal cortex ^53^. Subsequent analyses were conducted using custom software written in MATLAB (Mathworks). Fluorescent intensities of the pixels within an ROI were averaged to represent the signal of the ROI, denoted *F*, and this value was divided by the baseline signal value *F0*, which was calculated as an average of *F* across all frames, to obtain normalized fluorescence changes *dF/F* = (*F-F0)/F0*.

The extent of fluorescence signals derived from intrinsic sources (flavin fluorescence and hemodynamics ^56–58^) was estimated via the following two control experiments: imaging non-GCaMP6-expressing C57BL/6 (non-G6) mice (Figure S2) and correction of hemodynamic signals using two-wavelength imaging (Figure S5). In the former approach, basal fluorescence images of the dorsal cortical surface were acquired from non-G6 mice in order to estimate a potential upper bound of signal contamination. The average baseline signal intensity of non-G6 mice across three representative ROIs (M2a, HLp, and V1a) was 41.1 ± 0.9 % of that in Emx1G6 mice (Figure S2B, mean ± SEM, *n* = 7 Emx1G6 mice and 3 non-G6 mice). Furthermore, the average fluorescence changes of non-G6 mice across all hemispheric ROIs after locomotion onset was 19.4 ± 2.2 % of Emx1G6 mice (mean ± SEM, Figures S2C and S2D). These results imply that intrinsic fluorescence signals are much weaker and less dynamic than GCaMP fluorescence and that they constitute at most ∼8 % of signal changes observed in Emx1G6 mice. These observations are consistent with other recent studies ^59,60^ that were conducted without compensation of endogenous signals.

In the latter approach, we imaged Ca^2+^-dependent and -independent fluorescence signals at 470 and 405 nm wavelengths, respectively, in a separate cohort of Emx1G6 mice (*n* = 6), by following the previously described procedure ^61^. Images of fluorescence excited at 470 and 405 nm were captured alternately at an overall frame rate of 40 frames per second (20 frames per second for each wavelength) using two LED drivers (470 nm, SOLIS-470C and DC20; 405 nm, M405L4 and LEDD1B, Thorlabs) controlled by custom LabVIEW software. A hemodynamic correction was conducted by subtracting *dF/F*_*405 nm*_ from *dF/F*_*470 nm*_, where *dF/F*_*405 nm*_ and *dF/F*_*470 nm*_ are normalized fluorescence changes for signals obtained at 405 and 470 nm, respectively (Figure S5A; ^62^). The results demonstrate that although correlation coefficients between ROIs appeared slightly higher and thus resulted in identifying a larger number of highly correlated FCs (Figure S5B, see also Figure 2B), overall patterns of the network properties (node degree, betweenness centrality, CPL, and modularity Q, see below for details of these parameters) were qualitatively similar to those obtained without hemodynamic correction (Figures S5C–S5F, see also Figures 2C–2F; ^34^).

In this study, we focused on analyses of the dynamics of functional cortical networks during transitions between locomotion and rest. We considered only transitions from long rest to long locomotion (locomotion onset) and those from long locomotion to long rest (locomotion cessation). As a control, we randomly selected reference time points regardless of the behavioral state as many times as an average number of locomotion onset and locomotion cessation (random control). Sessions with at least two onset or two cessation events were included for analysis, and average fluorescence changes across all onset, cessation, or random events were calculated to obtain values representative of each session. The numbers of each type of transitions (onset, cessation, and random, respectively) analyzed are as follow: Emx1G6, 569, 653, and 659 events, *n* = 89 sessions from 7 mice; no projection, 382, 454, and 451 events, *n* = 71 sessions from 17 mice; hemodynamics correction, 658, 616, and 656 events, *n* = 71 sessions from 6 mice; Emx1G6_*15q dup*_, 275, 296, and 331 events, *n* = 88 sessions from 9 mice; Non-G6, 513, 621 and 590 events, *n* = 41 sessions from 3 mice.

To analyze functional connectivity (FC) between ROIs, we created correlation matrices representing the correlation between the cortical activity of all ROI pairs. We extracted 6-s segments of *dF/F* that spanned –3 s to +3 s around the event of interest (i.e., onset, cessation, or random). Each of the 6-s segments was then further divided into 6 non-overlapping 1-s subsegments, and FC was calculated as pair-wise Pearson correlation coefficients between *dF/F* during these 1-s subsegments. The 1-s time window was chosen to investigate rapid and dynamic changes of FC associated with behavior and accords with the recent notion that spontaneous behavior and ongoing brain activity are related to each other at a time scale of about 1 s ^63^. The correlation matrices obtained were averaged within a session and visualized as functional connectivity graphs of binarized networks using available MATLAB codes ^64^, in which the positions of ROIs were arranged according to their anatomical positions, and lines and symbol sizes represented highly correlated FC (*r* > 0.8) and the number of such connections associated with the ROI, respectively. The threshold for binarization (*r* > 0.8) selected top 26.7 ± 10.1 % of the most prominent connections out of all 1,225 connections between 50 ROIs (mean ± SD, *n* = 1,602 subsegments from 89 sessions times 3 conditions; average correlation coefficient, onset, 0.68 ± 0.03; cessation, 0.68 ± 0.01; random, 0.68 ± 0.01; mean ± SD, *n* = 534 subsegments from 89 sessions). Node degree, betweenness centrality, characteristic path length (CPL), and modularity Q were calculated using the Brain Connectivity Toolbox ^21^. Node degree and betweenness centrality represent the number of functional connections associated with each cortical ROI and the extent to which the ROI falls on the shortest paths between any other pairs of ROIs in the network, respectively. CPL represents the average shortest path length between all ROI pairs in the network. Modularity Q represents an index of optimized modules that maximize the number of within-module edges and minimize the number of between-module edges.

### Support vector machine classification

Support vector machine (SVM) classification was performed using the Statistics and Machine Learning Toolbox in MATLAB. For behavior state classification, we used the “fitclinear” function to train a linear classification model with high-dimensional predictor data. The SVM was regularized by the lasso method to reduce model complexity and prevent overfitting. The FC datasets included 17,970-time point data that spanned the entire 10-min sessions. Each time point data contained 1,225 FCs from 50 ROIs as features. The FCs were calculated using a 1-frame sliding window of 30-frame size without excluding short locomotion and short rest periods. The corresponding behavioral data were binary vectors in which rest and locomotion were labeled as 0 and 1, respectively. Data that contained at least two episodes of long locomotion or long rest within a session were used. All relevant data (11–15 sessions per mouse) from each mouse were concatenated to be used for training and testing. An SVM classifier was trained using datasets from four of all seven mice (train set), and binary classification was conducted on each time point of the FC data from the remaining three mice (test set). All 35 combinations arising from seven mice (_7_*C*_4_) were tested. Accuracy was calculated as a percentage of time points that were classified to the right behavioral state. The chance level was defined as the overall average percentage of short and long locomotion periods (58 %) since SVM tends to classify data to the more frequent category. In control, the datasets used for training were also used for testing (“train” control). In shuffled control, elements of behavioral state vectors were randomly shuffled and used for training and testing.

To identify features that contributed to the classification, we sorted features of the trained classifiers by their weights that represented coefficients of normal vector on the hyperplane. We then counted the appearance of features that included each ROI in the top 0.5 % and bottom 0.5 % distributions (6 features each, 12 total) as an importance index for the ROI. When cortical areas of interest contained multiple ROIs, this index was normalized by their number. The change level was defined as an average of 100 times random sampling. We then newly trained SVM classifiers using these top 0.5 % and bottom 0.5 % features and classified the test datasets to confirm that the selected features contribute to the classification. As a control, we tested classifiers trained with the same number of randomly selected features. This was repeated 100 times, and the results were averaged.

For genotype classification, we used the “fitcsvm” function in MATLAB, which trains and cross-validates an SVM model to solve problems with low-dimensional predictors. The 1,225 correlation coefficients were averaged throughout a session at each time point within relevant behavioral states or transitions. The data were then concatenated together for all relevant sessions, and the correlation coefficients were normalized into z-scores. We then cross-validated the classifiers using the leave-one-subject-out (LOSO) method, in which a pair of datasets from a mouse per each genotype were excluded from training and used for testing. In training, the classifiers were subjected to 10-fold cross-validation. All 63 combinations were tested from nine Emx1G6_*15q dup*_ mice and seven Emx1G6 mice.

### Histology

Mice were deeply anesthetized with isoflurane and perfused transcardially with phosphate-buffered saline (PBS) followed by 4 % paraformaldehyde (PFA) in PBS. Brains were removed and further fixed in 4 % PFA in PBS at 4°C overnight. Frozen parasagittal sections were cut on a cryostat to a thickness of 30 μm. The sections were incubated at 4°C overnight with rabbit anti-GFP antibody (1:1000, A-11122, Thermo Fisher) and mouse anti-GAD67 antibody (1:1000 clone 1G10.2, Millipore) diluted in PBS containing 5 % normal goat serum and 0.3 % Triton X-100, followed by Alexa Fluor 488-or Alexa 568-labeled goat anti-rabbit or antimouse IgG antibody (1:500, A-11034 or A-11019, ThermoFisher) diluted in the same buffer at room temperature for 1 h. Cell nuclei were counterstained using VectaShield Mounting Medium with DAPI (Vector Laboratories). Fluorescence images were acquired using a Keyence BZ-9000 epifluorescence microscope equipped with a 4× or 10× objective.

### Statistics

To statistically test functional network connectivity, we used Network Based Statistic (NBS) Toolbox in MATLAB ^23^. NBS nonparametrically calculates familywise error rate-corrected *P*-values with 5,000 times permutation testing. Inter-areal activity in test conditions was considered significantly correlated or decorrelated if the correlation coefficient during the behavioral transitions was higher or lower than random control with *P* < 0.01. In comparison between Emx1G6 mice and Emx1G6_*15q dup*_ mice, the differences were considered significant when *P* < 0.05. Other statistical tests were performed using MATLAB or R. For two-group comparisons, Welch’s t-test was used when normal distributions were assumed. Otherwise, Wilcoxon rank-sum test was used. For comparisons between more than two groups, Wilcoxon rank-sum test with Bonferroni correction, one-way ANOVA and two-way ANOVA with Tukey-Kramer test were used.

## Acknowledgment

We thank Norihiro Sadato, Masaki Fukunaga, and Takahiko Koike for advice on the network analyses, Yasunori Hayashi for general support and Hiromu Monai for comments on this manuscript.

## Funding

The KAKENHI from JSPS: 19H04942 (NN), 17H05985 and 19H04942 (MS), and 16H06316, 16H06463, 16H06276, 21H00202, 21H04813 and 21K19351 (TT)

The Incentive Research Project grant from RIKEN (NN)

Japan Agency for Medical Research and Development (JP21wm0425011) (TT)

Japan Science and Technology Agency (JPMJMS2299) (TT)

Intramural Research Grant (30-9) for Neurological and Psychiatric Disorders of NCNP (TT)

The Takeda Science Foundation, Smoking Research Foundation, Tokyo Biochemical Research Foundation, Research Foundation for Opto-Science and Technology, Taiju Life Social Welfare Foundation, The Naito Foundation, and The Tokumori Yasumoto Memorial Trust for Researches on Tuberous Sclerosis Complex and Related Rare Neurological Diseases (TT)

## Author contributions

Conceptualization: NN

Methodology: NN, MS

Investigation: NN, YS, XF

Visualization: NN Supervision: OY, MS, JN, TT

Writing—original draft: NN

Writing—review & editing: MS, AZ, TT

## Data and materials availability

All data, code, and materials are made available by the authors upon reasonable request.

## Supplementary information

Figure S1. Validation of regions of interest by sensory and motor mapping.

Figure S2. Estimation of the contribution of intrinsic fluorescence signals to the total signals acquired from GCaMP transgenic mice.

Figure S3. Hierarchical clustering of cortical activity during locomotion onset and cessation.

Figure S4. Relationship between node degree, betweenness centrality, and fluorescence changes.

Figure S5. The functional cortical network after hemodynamic correction.

Figure S6. Fluorescent signal intensities in the cortical areas of Emx1G6 and Emx1G6_*15q dup*_ mice.

Figure S7. Sensory mapping of Emx1G6_*15q dup*_ mice.

Figure S8. Abnormal correlations and decorrelations among cortical areas during behavioral transitions in Emx1G6_*15q dup*_ mice.

