## Supplementary information for "VR-based real-time imaging reveals abnormal cortical dynamics during behavioral transitions in a mouse model of autism"

**This PDF file includes:**

Figures S1 to S8

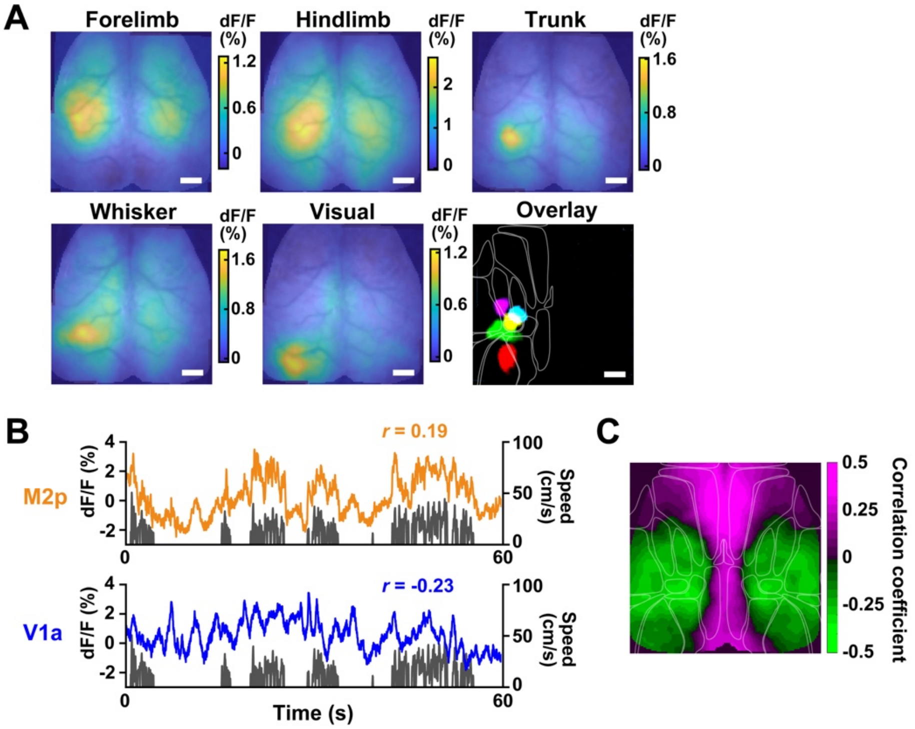


**Figure S1. Validation of regions of interest by sensory and motor mapping.**
(**A**) Validation of sensory cortical regions of interest (ROIs) by sensory mapping. Cortical areas that responded to tactile stimuli to the right forelimb, hindlimb, trunk, and whiskers and visual stimulus to the right eye are shown. An overlay of these five areas is shown at the bottom right. Scale bar = 1 mm.
(**B**) Example traces of fluorescence intensity in the posterior secondary motor cortex (M2p, top) and the anterior primary visual cortex (V1a, bottom). The black traces indicate the locomotion speed of the mouse during the corresponding period of time. The *r* indicates the correlation coefficient between fluorescence intensity and locomotion speed.
(**C**) Validation of motor cortical ROIs by pixel-based motor mapping. Pixels at which the fluorescence was positively and negatively correlated with locomotion speed are indicated in magenta and green, respectively.

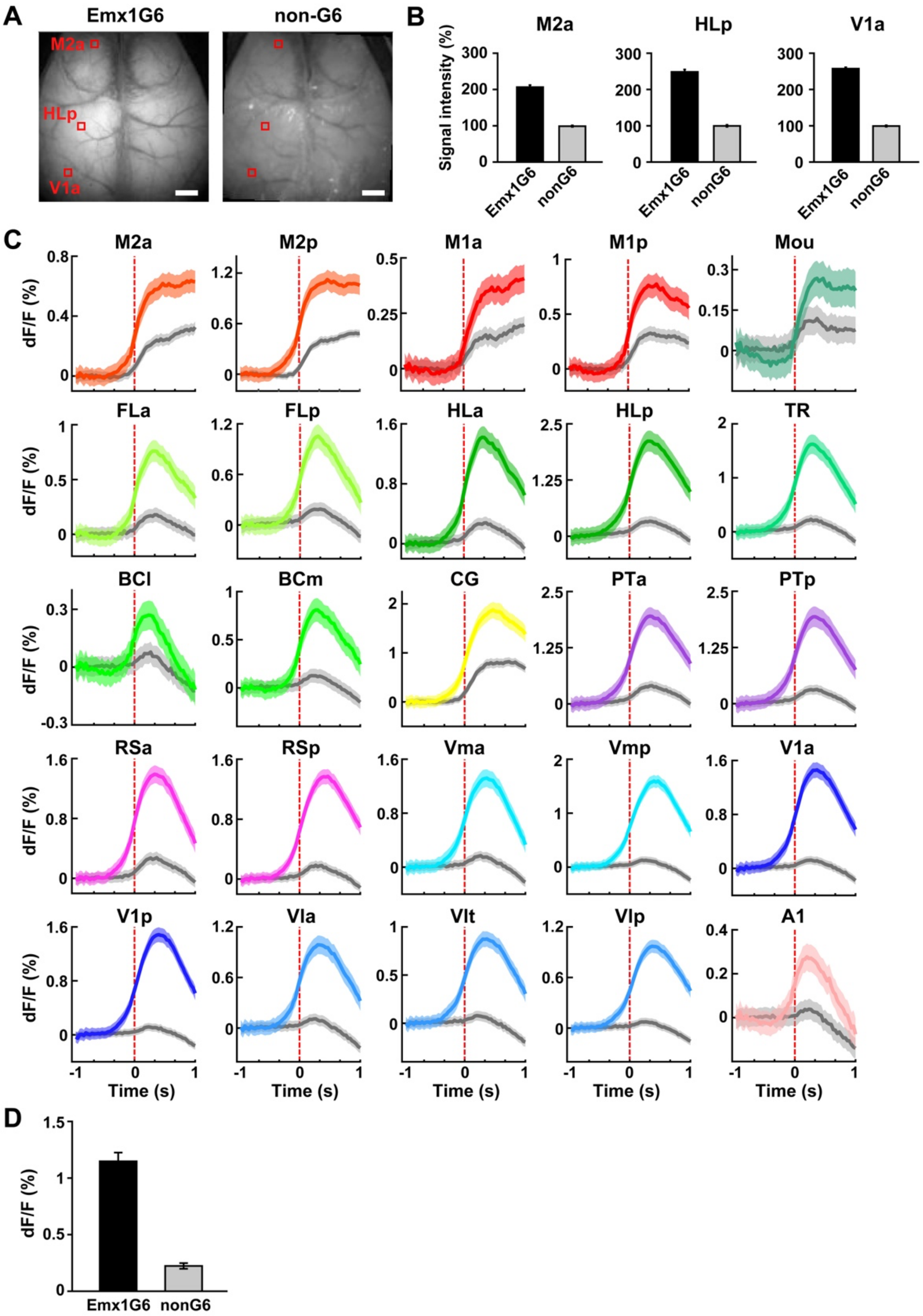


**Figure S2. Estimation of the contribution of intrinsic fluorescence signals to the total signals acquired from GCaMP transgenic mice.**

(**A**) Basal fluorescence images of the dorsal cortex acquired from Emx1G6 mice and C57BL/6 mice (non-G6). The red squares defined in M2a, HLp, and V1a represent ROIs for quantification in B. Scale bar = 1 mm.

(**B**) Fluorescence intensities quantified at M2a, HLp, and V1a. Data represent mean ± SEM ($n$ = 7 Emx1G6 mice and 3 non-G6 mice).
(**C**) Fluorescence changes around locomotion onset in Emx1G6 (colored) and non-G6 (gray) mice. Data for all 25 ROIs from the left hemisphere are shown (mean ± SEM, $n$ = 7 Emx1G6 mice and 3 non-G6 mice). Average values from −1 s to −0.67 s were used as a baseline.
(**D**) Average fluorescence changes across all ROIs. Data represent mean ± SEM ($n$ = 7 Emx1G6 mice and 3 non-G6 mice).

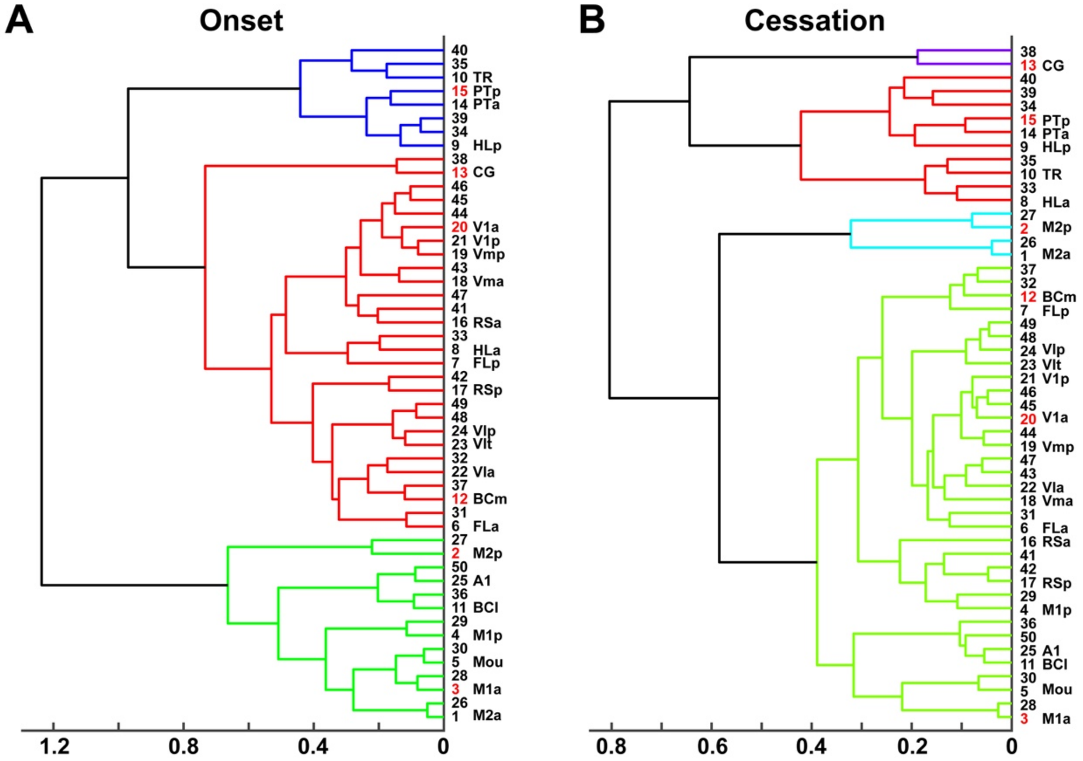


**Figure S3. Hierarchical clustering of cortical activity during locomotion onset and cessation.**

(**A**) Clustering of cortical activity during locomotion onset. Numbers on the right-hand side indicate ROI numbers, and fluorescence traces of ROIs shown by red numbers are presented in Figure 2A. Only ROI numbers for the left hemisphere are labeled with the corresponding cortical areas. The X-axis represents the height of the nodes (i.e., dissimilarity) expressed in Chebyshev distance. Clusters with less than 70 % of maximum dissimilarity were shown in the same colors. The same convention applies to (B).

(**B**) Clustering of cortical activity during locomotion cessation.

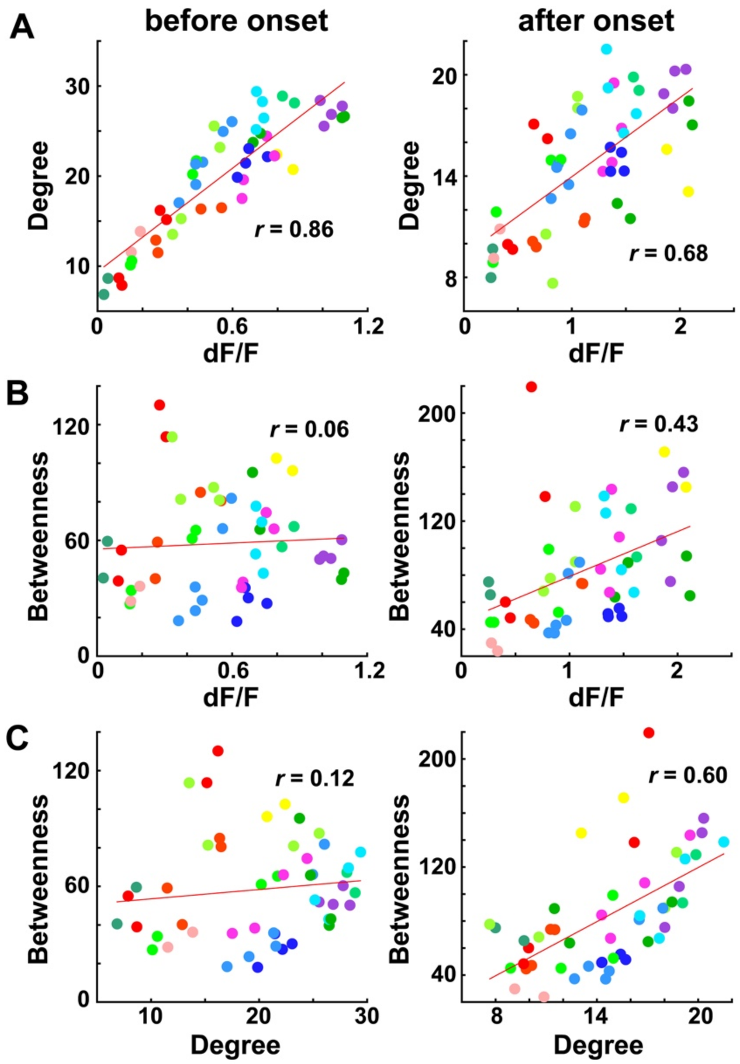


**Figure S4. Relationship between node degree, betweenness centrality, and fluorescence changes.**

(**A**) Scatter plots showing the relationship between fluorescence change (dF/F) and node degree for all 50 ROIs during a 1-s time window immediately before (left) and after (right) the locomotion onset. The *r* indicates the correlation coefficient between dF/F and node degree. Response amplitude of each ROI was significantly correlated with node degree both before and after the onset.

(**B**) Scatter plots showing the relationship between dF/F and betweenness centrality for all 50 ROIs. Response amplitude of each ROI was not correlated with betweenness centrality before the onset and weakly correlated after the onset.

(**C**) Scatter plots showing the relationship between node degree and betweenness centrality for all 50 ROIs. Node degree and betweenness centrality were not correlated before the onset, whereas they were weakly correlated after the onset.

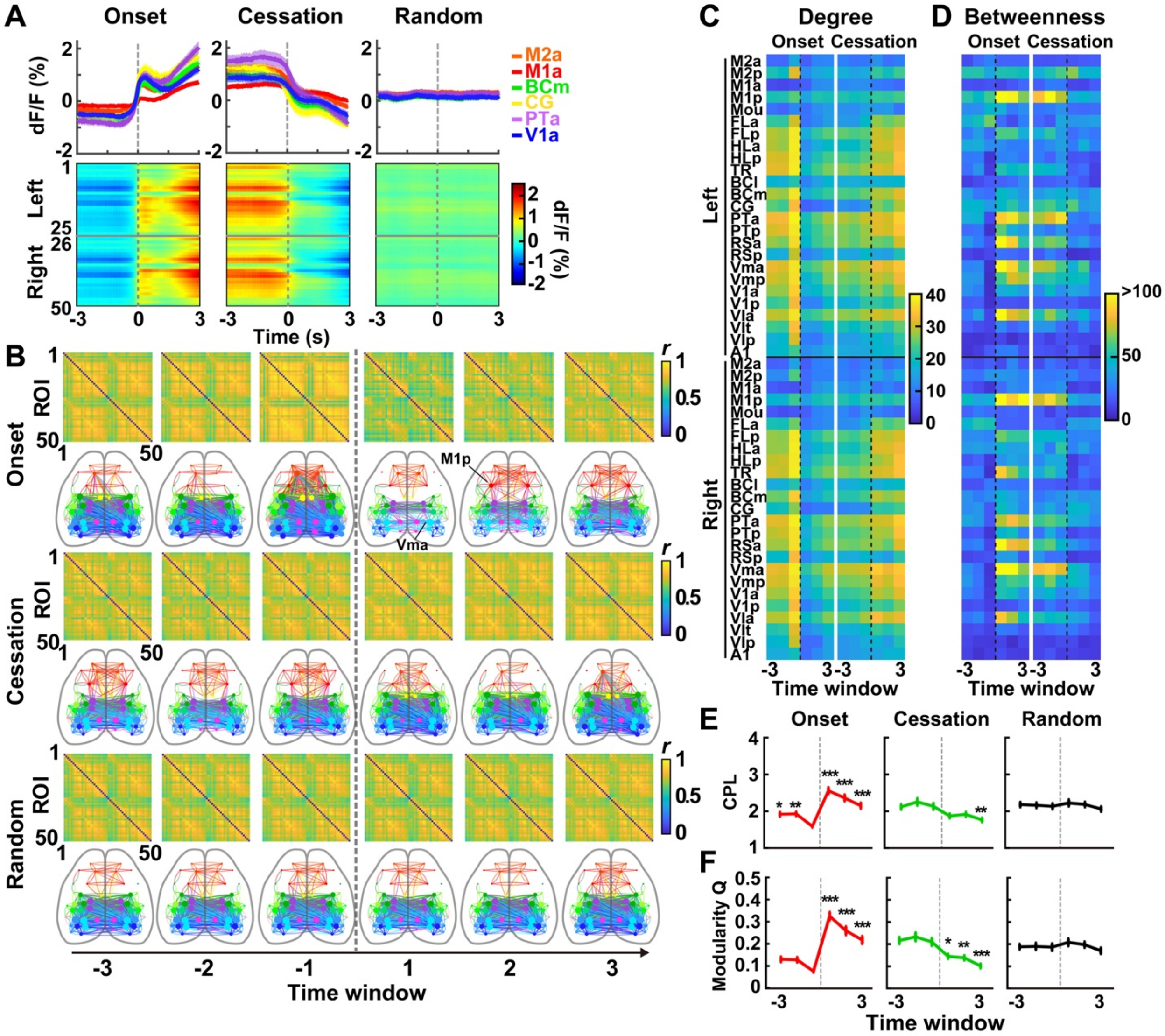


**Figure S5. The functional cortical network after hemodynamic correction.**

(**A**) Cortical activity of Emx1G6 mice after hemodynamic correction. The top plots present average relative changes in activity signals in representative cortical areas at locomotion onset, locomotion cessation, and random control ($n$ = 71 sessions from 6 mice). The colormaps at the bottom indicate changes in activity signals in all ROIs sorted by ROI number. The convention of the figure is the same as Figure 2A.

(**B**) Dynamics of correlations between activities of ROI pairs during the behavioral transition after hemodynamic correction. Correlation matrices and functional connectivity graphs (FC, $r > 0.8$) were generated using the data shown in (A). The convention of the figure is the same as Figure 2B.

(**C, D**) Changes in node degree (C) and betweenness centrality (D) during the behavioral transitions in each ROI of Emx1G6 mice after hemodynamic correction.

(**E**) Change in characteristic path length (CPL) during the transitions after hemodynamic correction. Data represent mean ± SEM. Onset: $F_{(5, 414)} = 42.33$, $P = 3.4\times10^{--35}$; Cessation: $F_{(5, 414)} = 13.92$, $P = 1.3\times10^{-12}$; Random: $F_{(5, 420)} = 1.44$, $P = 0.21$. *$P < 0.05$, **$P < 0.01$, ***$P < 0.001$, vs. time window -1, one-way ANOVA with Tukey Kramer test. $n$ = 71 sessions from 6 mice.

(**F**) Change in modularity Q during the transitions after hemodynamic correction. Data represent mean ± SEM., Onset: $F_{(5,\ 414)} = 25.5$, $P = 1.9\times10^{-22}$; Cessation: $F_{(5,\ 414)} = 6.69$, $P = 5.2\times10^{-6}$; Random: $F_{(5,\ 420)} = 1.38$, $P = 0.23$. *$P < 0.05$, **$P < 0.01$, ***$P < 0.001$, vs. time window -1, one-way ANOVA with Tukey Kramer test. $n = 71$ sessions from 6 mice.

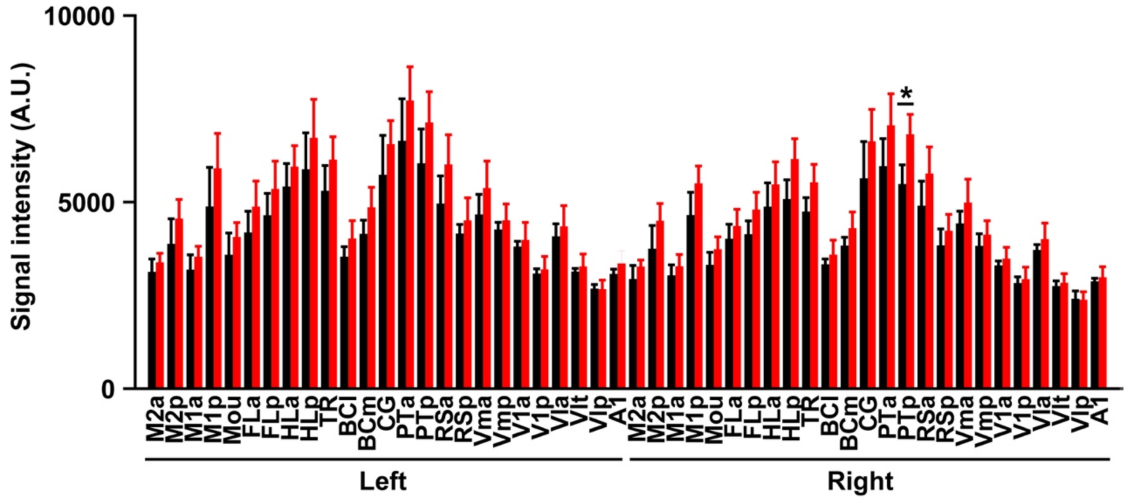


**Figure S6. Fluorescence signal intensities in the cortical areas of Emx1G6 and Emx1G6$_{15q\ dup}$ mice.**

(**a**) Average signal intensity was shown for each ROI in Emx1G6 mice (black) and Emx1G6$_{15q\ dup}$ mice (red). Data indicate mean ± SD. * $P$<0.05, t-test with Bonferroni correction. $n$ = 7 Emx1G6 mice and 9 Emx1G6$_{15q\ dup}$ mice.

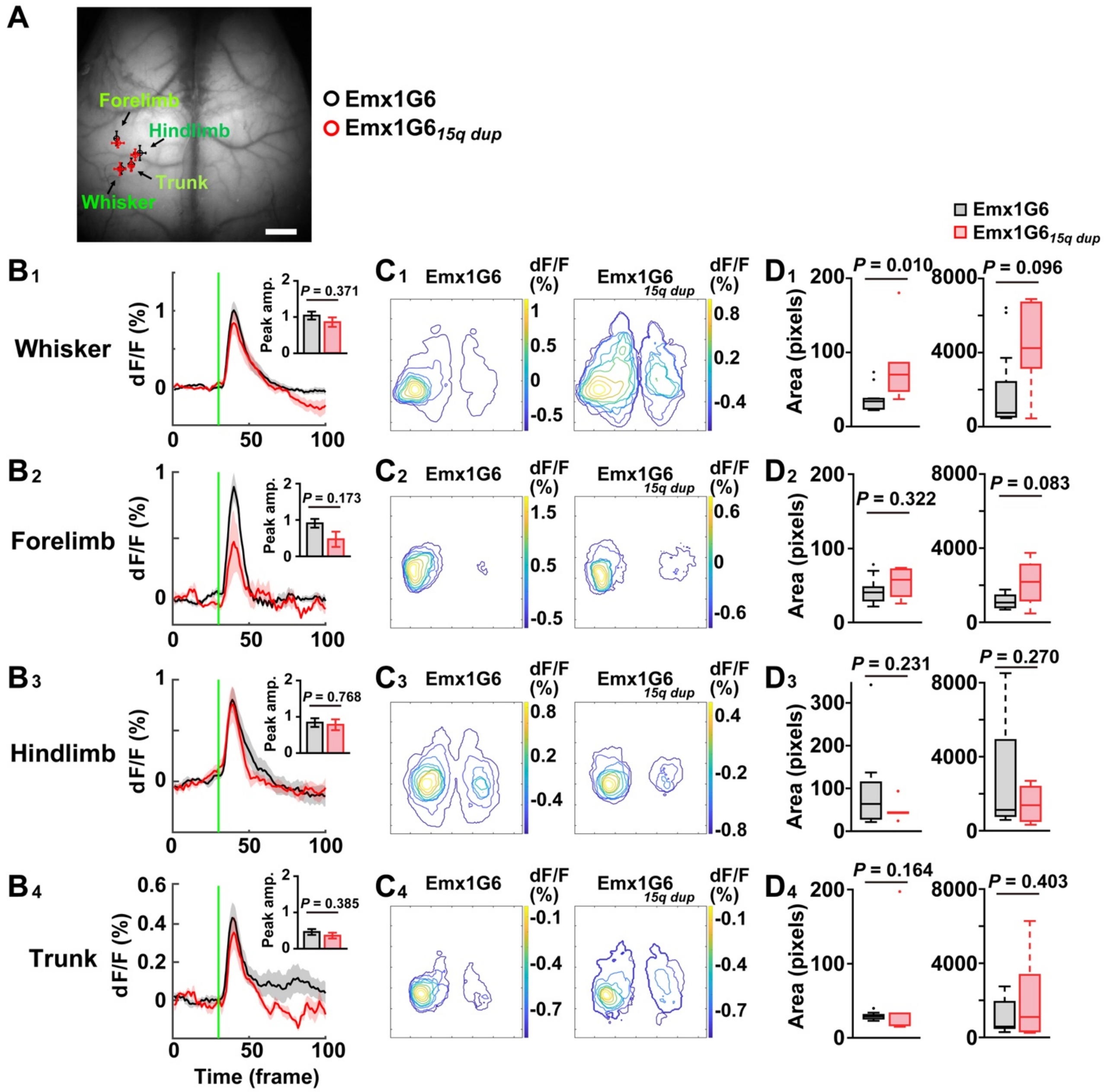


**Figure S7. Sensory mapping of Emx1G6$_{15q\ dup}$ mice.**

(**A**) The coordinates of peak responses to tactile stimulation to right whiskers, forelimb, hindlimb, and trunk in Emx1G6 mice and Emx1G6$_{15q\ dup}$ mice. The circles and error bars indicate mean ± SEM. Scale bar, 1 mm.

(**B-E**) Tactile responses in Emx1G6 mice and Emx1G6$_{15q\ dup}$ mice. (B$_1$-B$_4$) Fluorescence changes in response to tactile stimuli to right whiskers (B$_1$), forelimb (B$_2$), hindlimb (B$_3$), and trunk (B$_4$). Green lines indicate the delivery of tactile stimulation. Solid lines and shades indicate mean ± SEM (Whiskers, $n$ = 11 Emx1G6 and 5 Emx1G6$_{15q\ dup}$ mice; Forelimb, $n$ = 9 Emx1G6 and 4 Emx1G6$_{15q\ dup}$ mice; Hindlimb, $n$ = 10 Emx1G6 and 5 Emx1G6$_{15q\ dup}$ mice; Trunk, $n$ = 10 Emx1G6 and 5 Emx1G6$_{15q\ dup}$ mice). Inset bar graphs indicate peak response amplitudes. $P$ values in the t-test are shown. ($C_1$-$C_4$) Response areas. The contours in different colors indicate areas of 10–90 % peak responses at 10 % intervals. ($D_1$-$D_4$) Areas of > 90 % peak amplitude (left) and > 60 % peak amplitude (right). The boxes represent the 25th, 50th, and 75th percentiles, and the whiskers represent the range except for outliers. *P* values in the Wilcoxon rank-sum test are shown.

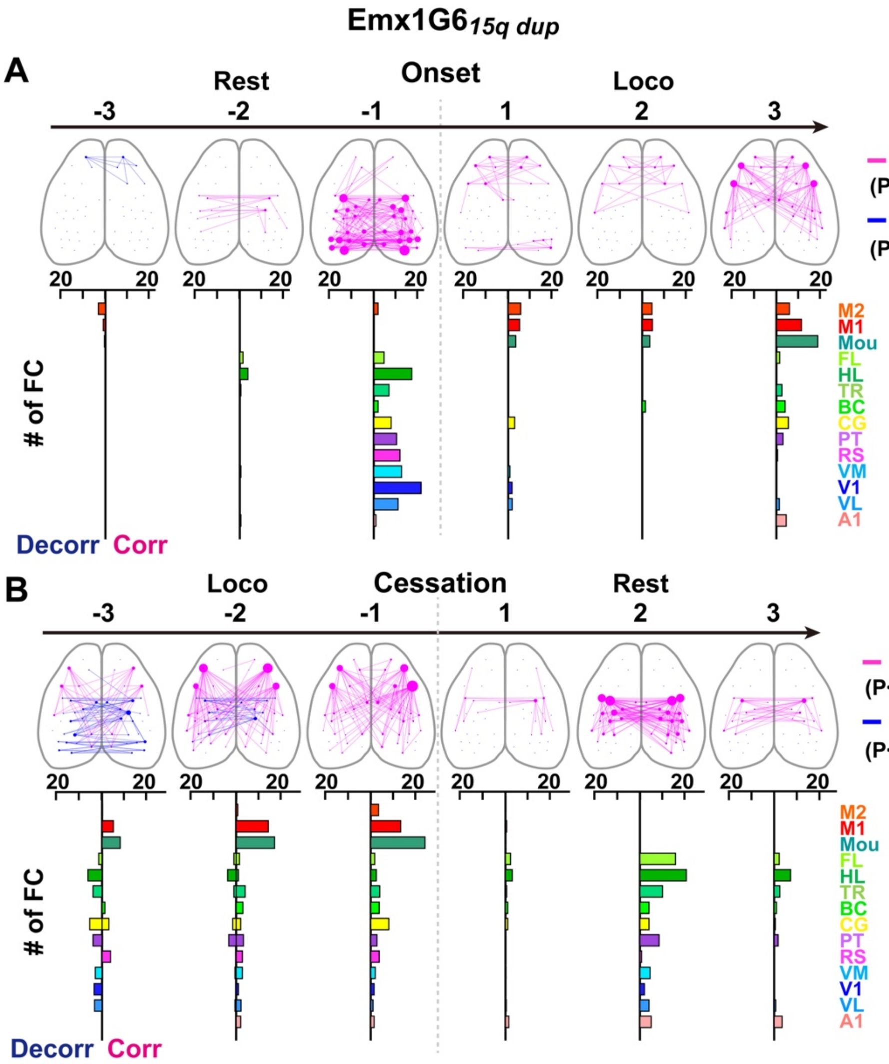


118 **Figure S8. Abnormal correlations and decorrelations among cortical areas during behavioral transitions in Emx1G6*15q dup* mice.**

120 (**A, B**) Significant correlations and decorrelations of functional cortical subnetworks of Emx1G6*15q dup* mice during locomotion onset (A) and cessation (B). *P*<0.01, NBS. See Figure 4A legend for figure convention.
